# Functional characterization of multi-domain LPMOs from marine *Vibrio* species reveals modulation of enzyme activity by domain-domain interactions

**DOI:** 10.1101/2025.07.04.663128

**Authors:** Yong Zhou, Eirik G. Kommedal, Zarah Forsberg, Gustav Vaaje-Kolstad, Wipa Suginta, Vincent G. H. Eijsink

## Abstract

Several bacterial pathogens secrete multi-domain enzymes known as lytic polysaccharide monooxygenases (LPMOs) that are important for virulence. One example is the *Vibrio cholerae* virulence factor GbpA (*Vc*GbpA), in which an N-terminal LPMO domain is followed by two domains of unknown function called GbpA2 and GbpA3, and a C-terminal chitin-binding domain, called CBM73. In-depth functional characterization of full-length and truncated variants of *Vc*GbpA and a homologue from *V. campbellii* (previously *V. harveyi*; *Vh*GbpA) showed that the catalytic LPMO domains of these proteins exhibit properties similar to natural single-domain LPMOs with established roles in chitin degradation. Interestingly, binding to chitin and efficient degradation of this substrate were affected by the presence of the GbpA2 and GbpA3 domains. Combined with structural predictions and analyses of sequence conservation, our data suggests that GbpA3 has evolved to interact with the reduced copper site to prevent off-pathway reactions in the absence of substrate. Substrate binding by CBM73 weakens this interaction, enabling the activation of the LPMO only when substrate is present. These observations shed new light into the functionality of these multi-domain LPMOs and uncover a novel mechanism for regulating LPMO activity.

## 1. Introduction

Chitin is a natural amino polysaccharide in which two *N*-acetyl-D-glucosamine (GlcNAc) sugars linked by a β-(1,4) glycosidic bond and rotated 180° relative to each other make up chitobiose, the repeating unit of the chitin polymer (1–3). Single chitin chains pack together to form a crystalline polysaccharide microfibril that is insoluble in water and confers mechanical stability to fungal cell walls and the exoskeletons of crustaceans and insects (2). Chitin may occur in complex co-polymeric structures, such as fungal cell walls, which also contain glucans, mannans, and glycoproteins (4, 5). Enzymatic conversion of chitin is of interest for many reasons: chitin is an abundant bioresource that can be valorized (6); chitin turnover is important for the global carbon cycle, especially in aquatic ecosystems (7); and chitin-degrading enzyme systems are associated with a wide variety of host-pathogen interactions and with microbial competition (8–13). Some microorganisms have evolved complex enzymatic systems to depolymerize chitin to GlcNAc, which can be utilized as a nutrient source. Next to endo- and exo-acting glycoside hydrolases (chitinases), these systems include lytic polysaccharide monooxygenases (LPMOs), which catalyze oxidative cleavage of glycosidic bonds in crystalline chitin microfibrils, thus providing new access points for further depolymerization by hydrolytic enzymes (14–18).

LPMOs (EC 1.14.99.53-56) are monocopper enzymes that utilize hydrogen peroxide (H_2_O_2_) to oxidize the glycosidic bonds of chitin and other crystalline polysaccharides (15, 19–22). Since the discovery of their catalytic activity in 2010 (15), LPMOs have received considerable attention due to their intriguing catalytic mechanism (21–26) and their importance for industrial bio-refineries (27–29). The CAZy database currently contains eight sequence-based LPMO families, which are referred to as Auxiliary Activities (AA), with family AA10 containing bacterial chitin-active LPMOs. The single catalytic copper ion of LPMOs is coordinated by a conserved histidine brace (19) and copper reactivity is modulated by nearby amino acids in what is called the second coordination sphere (30, 31). LPMOs were originally considered monooxygenases (R-H + O_2_ + 2 e^-^ + 2H^+^ ➔ R-OH + 2 H_2_O) employing O_2_ as co-substrate (15, 19, 20). Recent works have demonstrated that LPMOs operate as peroxygenases (R-H + H_2_O_2_ ➔ R-OH + H_2_O) using H_2_O_2_ as co-substrate (21, 22, 32–36). In both reactions, the LPMO needs to be reduced by a reductant; in the monooxygenase reaction, further supply of electrons and protons is required during catalysis, whereas these are delivered in the form of H_2_O_2_ in the peroxygenase reaction. Importantly, for convenience, LPMO reactions are frequently run under apparent monooxygenase conditions (LPMO + substrate + reductant, aerobic), i.e., without exogenously added H_2_O_2_. In such “reductant-driven” reactions, LPMO activity is limited by the rate of *in situ* H_2_O_2_ generation resulting from LPMO-catalyzed or abiotic oxidation of the reductant (37–40). These reductant-driven reactions are usually excessively slow, while reactions supplied with H_2_O_2_ are orders of magnitude faster (32–36).

Importantly, high concentrations of H_2_O_2_, which may emerge as a result of both *in situ* production or too high levels of H_2_O_2_ feeding, may lead to autocatalytic enzyme inactivation, especially at a low substrate concentration (21, 41). A considerable fraction of LPMOs contain a carbohydrate-binding module (CBM) that promotes substrate binding. By doing so, CBMs increase the chance that available H_2_O_2_ is used productively, i.e., to cleave the substrate through a peroxygenase reaction, rather than non-productively, in a futile peroxidase reaction that may lead to enzyme damage (21, 42, 43). It has been pointed out that, in reactions driven by ascorbate oxidation, enzyme inactivation may be a self-reinforcing process (44): as the LPMO becomes damaged, copper is released (45), promoting abiotic ascorbate oxidation and further increasing H_2_O_2_ levels, thereby accelerating enzyme inactivation.

In recent years, several multi-modular LPMOs have been described (12, 46–52). Some of these multi-modular LPMOs contain both a CBM and a glycoside hydrolase domain with similar substrate specificity as the LPMO domain, and their role in biomass conversion thus seems obvious (47, 48). On the other hand, intriguingly, there is a large group of bacterial LPMOs that comprise one or two domains of unknown function in between an N-terminal chitin-active LPMO domain and a C-terminal chitin-binding CBM, most often a CBM5/12 or CBM73. Both expression studies and the ecological niches in which these enzymes are found do not necessarily point to a role of these LPMOs in chitin conversion (49), and one may wonder if chitin is the biologically relevant substrate. Moreover, these LPMOs are often found in bacterial pathogens, and for several of these, a clear connection with virulence has been established (10, 12).

One example of these enzymes is the four-domain GbpA from *Vibrio cholerae* (*Vc*GbpA), which is a known virulence factor that was originally referred to as a GlcNAc-binding protein A, hence the abbreviation GbpA (11, 46). *Vc*GbpA consists of a chitin-active LPMO domain (53), two internal domains referred to as GbpA2 and GbpA3, and a C-terminal CBM73 domain with known affinity for chitin (54). The chitin-active LPMO domain is highly similar to other known chitin-active LPMOs such as *Sm*AA10A (originally known as CBP21), the chitin-active LPMO from *S. marcescens* (15). Recently, the complete structure of a homologous four-domain protein from *V. campbellii* (previously known as *V. harveyi*; *Vh*GbpA) was determined by X-ray crystallography (**Fig. 1A**), showing a rather elongated overall conformation, compatible with SAXS data for both *Vc*GbpA and *Vh*GbpA (46, 51). While the functional roles of the GbpA2 and GbpA3 remain unknown, it has been suggested that they may be involved in the interaction between *Vibrio* and relevant host surfaces (13, 55).

**Figure 1.**
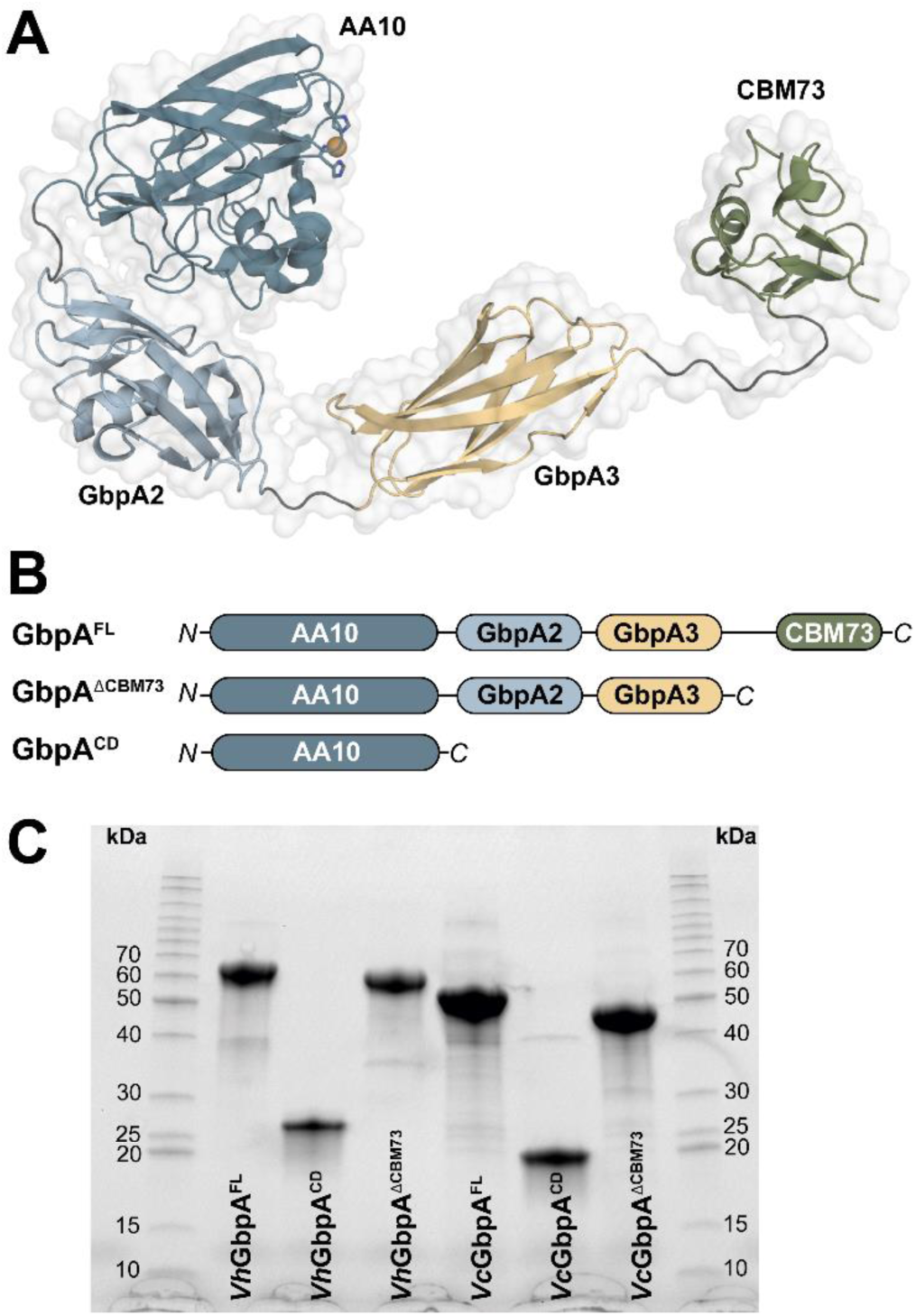
Overview of GbpA variants. (A) Crystal structure of full-length *Vibrio campbellii* GbpA (*Vh*GbpA^FL^) as determined by X-ray crystallography (PDB ID: 8GUL), illustrating the modular architecture of the protein; the catalytic copper appears as an orange sphere. (B) Schematic representation of the domain organization of full-length GbpA proteins and the two truncated variants generated in this study for both *V. cholerae* (*Vc*GbpA) and *V. campbellii* (*Vh*GbpA). (C) SDS-PAGE analysis of purified protein variants, i.e., full-length and truncated variants for *Vh*GbpA (left) and *Vc*GbpA (right).

To gain additional insight into the catalytic abilities of GbpA-like proteins and to investigate the roles of their individual domains, we carried out an in-depth functional characterization of full-length *Vh*GbpA, as well as two truncated variants: *Vh*GbpA^CD^, which contains only the catalytic LPMO domain, and *Vh*GbpA^𝛥CBM73^, which lacks the CBM73 domain. We studied multiple enzyme properties, such as binding to chitin, reduction and reoxidation rates, and the efficiency of on-and-off pathway reactions. Several of these experiments were also done with similar variants of *Vc*GbpA, showing similar results and adding confidence to the conclusions. Unexpectedly, the comparison of wild-type enzymes with truncated variants revealed an interaction between the combined GbpA2 and GbpA3 part of the protein and the catalytic domain. Structure predictions with AlphaFold3 (56) and sequence conservation patterns strongly suggest a copper-mediated interaction between GbpA3 and the catalytic copper site, which seems to have evolved to protect the enzyme from engaging in damaging off-pathway reactions. These observations lend support to a novel model in which *Vibrio* species regulate the activity of this virulence factor in a substrate-dependent manner.

## 2. Materials and methods

### 2.1 Materials

All chemicals were obtained from Sigma-Aldrich (St. Louis, MO, USA) unless specified otherwise. The model microcrystalline chitin substrate used in the study is commercial β-chitin from squid pen (5 mm flakes) acquired from Glentham Life Science Ltd. (Corsham, UK), which had been milled and sieved in-house to a particle size of < 75 μm using a PM200 planetary ball mill (Retsch, Haan, Germany) with zirconium oxide grinding tools. 100 mM ascorbic acid stock solutions were prepared in metal-free TraceSELECT water (Honeywell, Charlotte, NC, USA), filter-sterilized using 0.22 µm syringe filters, and aliquoted and stored at −20°C in light-protected tubes and were used only once after thawing. Tryptone and yeast extract were obtained from Thermo Fisher Scientific.

### 2.2 Protein production and purification

The gene coding for full-length *Vh*GbpA^FL^ (amino acid residues 24–487, AA10-GbpA2-GbpA3-CBM73, accession number ABU72648) was synthesized by GenScript Biotech (Piscataway, NJ, USA) as described previously (51), while the genes encoding the two truncated variants, the LPMO catalytic domain alone (*Vh*GbpA^CD^, amino acid residues 24–201) and the CBM73 truncated variant (*Vh*GbpA^𝛥CBM73^, amino acid residues 24–415) were synthesized after codon optimization for *E. coli* expression by Synbio Technologies (NJ, USA). The genes were cloned into pET-20b(+) for *Vh*GbpA^CD^ or pET-22b(+) for *Vh*GbpA^𝛥CBM73^, using NdeI/XhoI restriction sites, leading to in-frame fusions with nucleotide sequences encoding the 22-amino-acid *pelB* signal peptide at the N-terminus, to facilitate protein export to the periplasm, and six histidine residues at the C-terminus.

Wild-type *Vh*GbpA^FL^ was expressed and purified as described previously (51). To produce the truncated forms, the plasmids were transformed by heat shock into chemically competent One Shot^®^ BL21 Star^TM^ (DE3) cells (Invitrogen). Fresh colonies were inoculated in Luria-Bertani broth (LB medium) supplied with 100 µg.mL^-1^ ampicillin. To induce protein expression, the bacteria were grown in LB medium (37°C, 220 rpm) to an OD_600_ of 0.6∼0.8 before induction with isopropyl-β-D-thiogalactoside (IPTG) at a final concentration of 0.2 mM and then grown at 20°C, 200 rpm for 16 hours. Cells were harvested by centrifugation (8000 ×g for 10 mins at 4°C) using a Beckman Coulter centrifuge (Brea, CA, USA), after which periplasmic proteins were extracted using cold osmotic shock with 1.0 mM of magnesium chloride, as previously described (57). The resulting periplasmic fractions were sterilized by filtration through a 0.22-μm syringe filter prior to storage at 4°C and subsequent purification of the enzyme. The extracts were subjected to anion exchange chromatography using a 5-ml HiTrap HP (Q Sepharose) column (GE Healthcare, Chicago, USA). After the application of up to 30 mL sample, the column was washed with 10 column volumes of 20 mM Tris–HCl, pH 7.5, followed by elution with a linear gradient of NaCl (0–1 M over ten column volumes) in 20 mM Tris–HCl, pH 7.5, at a flow rate of 2 ml.min^-1^. LPMO-containing fractions were identified by SDS-PAGE, pooled, concentrated to approximately 2 mL using Vivaspin ultrafiltration tubes, 10 kDa MWCO (Sartorius, Göttingen, Germany) and loaded onto a ProteoSEC Dynamic 16/60 3–70 HR preparative size-exclusion column (Protein Ark, Sheffield, UK) operated at 1 mL.min^-1^ flow rate and equilibrated with 20 mM Tris-HCl, pH 7.5. The purity of resulting protein preparations was confirmed by SDS-PAGE. Protein concentrations were determined using UV spectroscopy at 280 nm and theoretical extinction coefficients, which were predicted using the Expasy ProtParam tool.

*Vc*GbpA^FL^ and its truncated variants, *Vc*GbpA^CD^ and *Vc*GbpA^𝛥CBM73^, were produced and purified as previously reported, without a His-tag on the C-terminus (46). The GH20 β-N-acetylhexosaminidase, or chitobiase, from *Serratia marcescens*, here referred to as *Sm*CHB, was recombinantly produced and purified as previously described (53). The GH18 chitinases A and C from *Serratia marcescens* (*Sm*Chi18A and *Sm*Chi18C) were recombinantly produced as described previously (58).

### 2.3 Copper saturation

Solutions with purified LPMOs (in 20 mM Tris-HCl, pH 7.5) were concentrated to reach a total volume of about 2 mL using Vivaspin ultrafiltration tubes (10 kDa MWCO; Sartorius, Göttingen, Germany), after which they were supplemented with a two-fold molar surplus of Cu(II)SO_4_, followed by incubation at room temperature for 30 mins. In some cases, this was done prior to the size-exclusion chromatography step of the purification protocol, which subsequently removed excess copper. To ensure the removal of free copper in protein samples treated after size exclusion chromatography, the samples were subjected to multiple consecutive rounds of concentration and dilution in 20 mM Tris-HCl, pH 7.5, using Amicon® Ultra-15 Centrifugal filter units (Merck, Dramstadt, Germany). The resulting total dilution factor for solutions of copper-saturated LPMOs amounted to at least 1,000,000-fold. Purified enzymes were stored at 4°C until use.

### 2.4 Reduction and reoxidation kinetics

Single turnover experiments to determine the rate of the reduction of LPMO-Cu(II) to LPMO-Cu(I) by ascorbate were performed using a single mixing set-up, while the reoxidation of LPMO-Cu(I) to LPMO-Cu(II) by H_2_O_2_ was assessed using a double-mixing set-up, essentially as described previously (59, 60), using the fluorescence change that accompanies reduction or oxidation of the LPMO for detection. All experiments were conducted with an SFM-4000 stopped-flow system equipped with a MOS-200M dual absorbance fluorescence spectrometer (BioLogic, Grenoble, France), with an applied voltage of 600 mV for detection. The excitation wavelength was set to 280 nm, and the increase (for reduction) or decay (for reoxidation) of fluorescence intensity was collected with a 340 nm bandpass filter. All experiments were carried out at 25 °C in 20 mM Tris-HCl, pH 7.5. Anaerobic conditions were obtained by storing N_2_–purged buffers and labware in a Whitley A95TG anaerobic workstation (Don Whitley Scientific, West-Yorkshire, UK) for 24 hours prior to preparing the necessary solutions in the anaerobic workstation. Purging with N_2_ was achieved using a Schlenk line, and all solutions were eventually transferred to sealed syringes in the anaerobic chamber.

To determine LPMO reduction rates, 75 μL of a 10 μM LPMO-Cu(II) solution (5 μM as final concentration) was mixed with 75 μL of an ascorbic acid solution with concentrations ranging from 50 to 800 μM (25-400 μM after mixing) at 25 °C, ensuring pseudo-first-order conditions ([E] < <[S]). For reoxidation, double-mixing experiments were performed in two steps. In the first step, the LPMO-Cu(II) (10 µM initial concentration) was mixed with one molar equivalent of L-cysteine for 10 s to form LPMO-Cu(I). In a second step, the *in-situ* generated LPMO-Cu(I) was mixed with different concentrations of H_2_O_2_ (ranging from 12.5 – 800 µM after mixing), followed by monitoring the decay of fluorescence intensity. The stopped-flow rapid spectrophotometer was flushed with a significant excess of deoxygenated buffer before connecting the anaerobically prepared syringes containing the solutions to be used and performing the experiments.

Pseudo-first order reaction rates (*k_obs_)* were determined by solving a single exponential equation with correction of baseline drift using the following equation: 𝑦 = 𝑎𝑡 + 𝑏 + 𝑐𝑒^−𝑘𝑜𝑏𝑠𝑡^. In all cases, plots of *k*_obs_ *vs.* [ascorbate] or [H_2_O_2_] were fitted using linear least squares regression to obtain the apparent second-order rate constant of the reduction or reoxidation step (*k*_AscA_ or *k*_H2O2_, respectively). All experiments were done at least in triplicates.

For *Vh*GbpA^FL^, reoxidation rates using the stopped-flow method could not be determined due to baseline drift of the reoxidized LPMO-Cu(II). Therefore, fluorescence spectroscopy was employed to investigate this reoxidation rate. The LPMO-Cu(II) (2 µM in a 1 mL fluorescence quartz cuvette) was initially mixed by stirring (sealed by parafilm M) with one molar equivalent of L-cysteine for 120 s to form LPMO-Cu(I), after which the generated LPMO-Cu(I) was mixed with different concentrations of H_2_O_2_ (ranging from 2 – 20 µM after mixing) followed by monitoring the decay of fluorescence intensity.

Alternatively, the reoxidation of LPMO-Cu(I) by O_2_ under ambient conditions (i.e., an O_2_ concentration of approximately 250 µM) was monitored by measuring the decay of fluorescence intensity, as previously reported, with minor modifications (60). 2 µM LPMO-Cu(II) was initially mixed with an equimolar amount of L-cysteine for 60 s to generate LPMO-Cu(I). The generated LPMO-Cu(I) was subsequently reoxidized by O_2_ in the solution. The decay of fluorescence intensity (excitation, 280 nm; emission, 342 nm) of LPMO-Cu(I) was recorded over the course of one hour and analyzed using a double exponential decay model with GraphPad Prism. Each experiment was performed in duplicates.

### 2.5 Substrate-binding assay

To examine the impact of the CBM73 domain on chitin-binding affinity, *Vh*GbpA variants [i.e., *Vh*GbpA^FL^ (10 μM), *Vh*GbpA^CD^ (20 μM), or *Vh*GbpA^𝛥CBM73^ (10 μM)] were incubated with 10 g.L^-1^ of β-chitin in 20 mM Tris-HCl (pH 7.5) at 22 °C and 1000 rpm for one hour, either in the absence or presence of 1 mM AscA, in a final volume of 200 µL. The binding time was counted from the moment ascorbate was added (or buffer, in the control without AscA), while a reaction without substrate was included as a control. The binding event was stopped by centrifuging at 22 °C and 13,000 rpm for 10 mins. The supernatant (up to 100 μL) was collected for further analysis. The pellet was resuspended and washed with 200 µL of the same buffer, and this washing step was repeated twice. The supernatant from the second-round washing was kept for analysis, while the pellet was resuspended in 200 µL SDS loading buffer and heated at 95 °C for 10 mins. All fractions were then analyzed by SDS-PAGE, with sample volumes adjusted to represent approximately the same proportion of the original reaction mixture.

Binding to β-chitin was also evaluated with the same time-course reaction conditions as for chitin degradation (see below). Reactions were performed at 22 °C using reaction mixtures containing 2 µM enzyme and 10 g.L^-1^ of β-chitin in 20 mM Tris-HCl, pH 7.5. Ascorbate was added to a final concentration of 1 mM and was the last reagent to be added. Samples were taken after 1, 3, 5, 10, 30, and 60 minutes of incubation and separated from the insoluble chitin by filtration using a 96-well filter plate (Merck) operated with a vacuum manifold (Millipore). The unbound protein in the filtrate was quantified with an adapted Bradford protocol for measuring low protein concentrations (61, 62). For each *Vh*GbpA variant, a control reaction without β-chitin was included and used as a reference to calculate the fraction (%) of unbound protein. Controls without enzymes were also added to monitor unspecific signals in the Bradford assay. All reactions were done in triplicates.

### 2.6 Reactions with chitin

Reductant-driven chitin-degradation experiments (“monooxygenase conditions”) were carried out in 2 mL Eppendorf tubes in an Eppendorf Thermomixer C (Eppendorf, Hamburg, Germany) set to 30 °C and 1 000 rpm. If not stated otherwise, 0.5 µM LPMO was pre-incubated with 10 g.L^-1^ squid pen β-chitin (Glentham Life Science Ltd., Corsham, UK) with a particle size of <75 μm for 30 mins at 30 °C before adding 1 mM AscA to start the reaction. All LPMO reactions were carried out in 20 mM Tris-HCl, pH 7.5.

The reactions were sampled at various time points, and enzyme activity in the samples was stopped by separating the LPMO from the insoluble substrate using a MultiScreen™ 96-well filter plate (Millipore, Burlington, MA, USA) operated with a Millipore vacuum manifold. Qualitative analysis of oxidized products was carried out by matrix-assisted laser desorption/ionization-time of flight mass spectrometry, MALDI-ToF MS, using a Bruker Autoflex mass spectrometer equipped with a 337 nm nitrogen laser. 1 μL of the reaction mixture was mixed with 1 μL of 2,5-dihydroxybenzonic acid (10 g·L^−1^ dissolved in 30% acetonitrile with 0.1% (v/v) trifluoroacetic acid) and applied onto an MTP 384 target plate ground steel TF (Bruker Daltonics GmbH, Bremen, Germany), followed by drying under a stream of air. Data acquisition and analysis were performed using FlexControl and FlexAnalysis (Bruker Daltonics GmbH).

For quantification of soluble oxidized chitooligosaccharides generated by the LPMO, the filtrates (35 μL) were transferred to new tubes and supplemented with *Sm*CHB to a final concentration of 0.5 μM, followed by static incubation at 37 °C overnight. Treatment with *Sm*CHB converts longer native and oxidized chitooligomers to a mixture of *N*-acetylglucosamine (GlcNAc, native monomer) and oxidized chitobiose (GlcNAcGlcNAc1A, oxidized dimer), which simplifies quantification of soluble reaction products, as described previously (53).

Reactions with exogenously added H_2_O_2_ were set up in the same manner. After the 30 min preincubation, varying concentrations of H_2_O_2_ (0, 50, 100 and 200 μM) were added after which the reaction was initiated by addition of 0.1 mM AscA. Sampling, sample treatment with *Sm*CHB and product analysis were performed as described above.

### 2.7 Quantification of reaction products

Quantification of chitobionic acid (GlcNAcGlcNAc1A) and *N*-acetylglucosamine (GlcNAc) was done by chromatography using an RSLC system (Dionex, Sunnyvale, CA, USA) equipped with a 100×7.8 mm Rezex RFQ-Fast Acid H^+^ (8 %) (Phenomenex, Torrance, CA, USA) column operated at 85 °C. Samples of 8 μL were injected into the column, and sugars were eluted isocratically for six minutes with 5 mM sulfuric acid as mobile phase and a flow rate of 1 mL.min^-1^. The analytes were monitored using a 194 nm UV detector. Standards of GlcNAcGlcNAc1A (25–800 μM) and GlcNAc (50–2500 μM) were used for quantification. GlcNAc was purchased from Megazyme (Bray, Ireland; 95 % purity), while GlcNAcGlcNAc1A was generated in-house by complete oxidation of *N*-acetyl-chitobiose (Megazyme, Bray, Ireland; 95 % purity) with a chitooligosaccharide oxidase from *Fusarium graminearum* (63), as previously described (53).

### 2.8 Oxidase activity

The oxidase activity of the various GbpA variants was measured using the protocol described by *Kittl et al.* (64) with modifications. Briefly, 50 µL of a reaction pre-mixture comprised of 0.2 mM Amplex Red and 10 U.mL^-1^ horseradish oxidase (HRP) in Tris-HCl buffer (20 mM, pH 7.5) was mixed with 40 µL LPMO (1 µM as the final concentration in the reaction) prepared in the same buffer and incubated at 30°C for 5 min. 10 µL of 10 mM AscA was added to start the reaction. The microtiter plate was shaken for 30 s at 600 RPM prior to recording absorbance at 563 nm every 30 s for 60 min. Control reactions without enzyme and with free copper instead of enzyme were included. The amount of H_2_O_2_ produced was quantified based on an H_2_O_2_ standard curve (0 to 20 µM), including 1 mM AscA. All reactions were performed in triplicates.

### 2.9 Determination of apparent melting temperature (*T*_m_)

The apparent melting temperature (*T*_m_) of *Vh*GbpA variants was determined using the protein thermal shift assay (Thermo Fisher Scientific, Waltham, MA, USA). This assay employs the fluorescent dye SYPRO Orange to monitor protein unfolding (65). The quantum yield of SYPRO orange increases significantly upon binding to hydrophobic regions that become exposed during protein unfolding. The fluorescence emission was monitored using a StepOnePlus real-time PCR machine (ThermoFisher Scientific). For the assay, 5 μM enzyme was prepared in 20 mM Tris-HCl buffer (pH 7.5), with or without 5 mM EDTA, and incubated with SYPRO Orange dye in a 96-well plate (MicroAmp® Fast Optical 96-well reaction plate with barcode 0.1 mL, Thermo Fisher Scientific) covered by and adhesive film (MicroAmpTM Optical Adhesive Film, Thermo Fisher Scientific). The temperature was gradually increased from 25 to 99°C over a duration of 75 minutes. Each experiment was carried out in triplicate (*n* = 3) to ensure reproducibility.

### 2.10 Ascorbic acid depletion

Reactions containing 0.5 µM of *Vh*GbpA variants and 1 mM ascorbic acid (AscA) were prepared by mixing equal volumes (50 µL each) of protein solution (1 µM) and AscA solution (2 mM) directly in a 96-well UV-transparent plate (Corning, Corning, NY, USA). AscA depletion was monitored spectrophotometrically by measuring absorbance at 255 nm every 3 min for up to 8 h using a Varioskan LUX plate reader (Thermo Fisher Scientific, Waltham, MA, USA). AscA concentrations were calculated using a standard curve generated from known AscA concentrations measured at 255 nm. All experiments were performed in triplicates (n = 3).

## 3. Results and Discussion

### 3.1 Protein production and assessment of redox properties

The two wild-type GbpA proteins and two similarly truncated variants of each were produced and purified (**Fig. 1**), yielding 4.5 to 25 mg of purified copper-saturated protein per liter of culture. Of note, in the below, the *Vc*GbpA and *Vh*GbpA variants generally show similar features despite the presence of a His-tag on the *Vh*GbpA variants, adding confidence to the results. For the *Vh*GbpA variants, protein integrity was assessed by determining apparent melting temperatures. All three proteins showed a clear unfolding transition with apparent melting temperatures between 45 and 50 ^°^C in reactions with EDTA and between 50 and 60 ^°^C in reactions without EDTA (**Fig. S1**). Copper binding is known to stabilize correctly folded LPMOs; hence, the observed destabilizing effect of EDTA was expected and indicates that the proteins do bind copper. Accordingly, all protein variants showed reductant-dependent activity on β-chitin as demonstrated by MALDI-ToF MS analysis of solubilized reaction products (**Figs. S2 & S3**).

The possible effects of the truncations on the reactivity of the copper site were assessed using a variety of methods. Measurements of the oxidase activity i.e., ascorbate-dependent production of H_2_O_2_ in the absence of substrate (44, 64), gave rates varying from 0.15 min^-1^ to 0.66 min^-1^ (**Table S1**), which are similar to the rates observed for other chitin-active bacterial LPMOs (44, 66). Interestingly, for both *Vh*GbpA and *Vc*GbpA, the full-length enzyme and the CBM73-truncated enzyme showed 2 -3 times lower oxidase rates than the isolated catalytic domain (see below for further discussion).

The second-order rates for reduction with ascorbic acid and reoxidation with H_2_O_2_ in the absence of substrate were determined using stopped-flow fluorimetry. This approach was based on observations by Bissaro *et al.* (59), who showed that the fluorescence signal of an LPMO depends on the redox state of the copper center (**Fig. S4**). Experiments were done using pseudo-first-order conditions using different concentrations of AscA in a single-mixing experiment to assess reduction, or H_2_O_2_, in a double-mixing experiment, to assess reoxidation (see Methods section for details). The total fluorescence change upon reduction was the same in all experiments, suggesting that reduction was complete in all cases. The reduction reaction remained monophasic but increased in rate with increasing concentrations of AscA. Likewise, the reoxidation reaction remained monophasic but increased in rate with rising H_2_O_2_ concentrations. Each fluorescence-versus-time curve was fitted to a single exponential function to derive pseudo-first-order rate constants (*k*_obs_), which were plotted as a function of [AscA] or [H_2_O_2_]. For both assays, the data could be fitted to a straight line, yielding the second-order rate constants *k*_AscA_ and *k*_H2O2_ (**Fig. S4**). The results (**Table 1**) show similar reduction (1.14 – 1.96 x 10^5^ M^-1^⋅s^-1^) and reoxidation (5.71 – 9.28 x 10^3^ M^-1^⋅s^-1^) rates for all protein variants. Although rate differences were small, it may be noted that the variants only comprising the catalytic domain showed the lowest rate of reduction and the highest rate of reoxidation for both *Vh*GbpA and *Vc*GbpA.

**Table 1.**
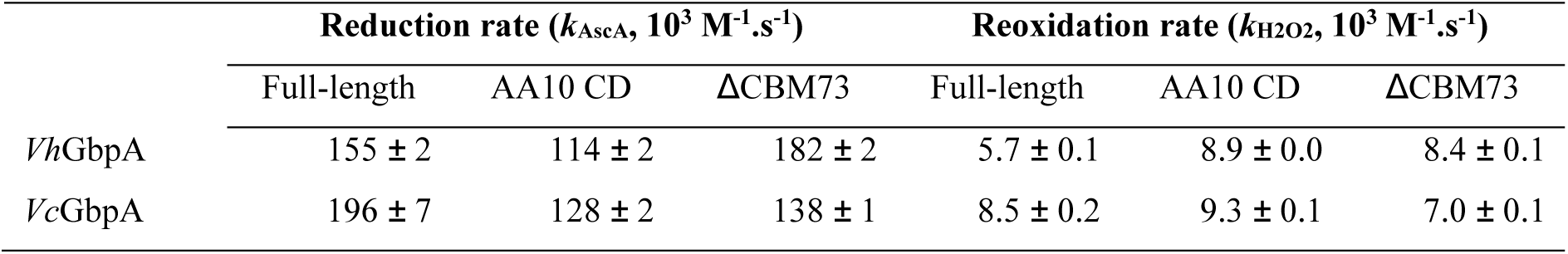
Reduction and reoxidation rates. The measurements were performed in 20 mM Tris-HCl, pH 7.5, with the underlying kinetic curves shown in **Fig. S4**. The reported values are derived from three independent experiments and are presented with standard deviations. Note that the method used for measuring reoxidation of *Vh*GbpA^FL^ deviates from the method used in all other cases; see Materials and Methods and **Fig. S4**.

| | Reduction rate ($k_{\text{AscA}}$, $10^3 \text{ M}^{-1}\cdot\text{s}^{-1}$) | | | Reoxidation rate ($k_{\text{H2O2}}$, $10^3 \text{ M}^{-1}\cdot\text{s}^{-1}$) | | |
| --- | --- | --- | --- | --- | --- | --- |
| | Full-length | AA10 CD | $\Delta\text{CBM73}$ | Full-length | AA10 CD | $\Delta\text{CBM73}$ |
| *VhGbpA* | $155 \pm 2$ | $114 \pm 2$ | $182 \pm 2$ | $5.7 \pm 0.1$ | $8.9 \pm 0.0$ | $8.4 \pm 0.1$ |
| *VcGbpA* | $196 \pm 7$ | $128 \pm 2$ | $138 \pm 1$ | $8.5 \pm 0.2$ | $9.3 \pm 0.1$ | $7.0 \pm 0.1$ |

The kinetic values derived from the stopped-flow experiments provide unprecedented insight into the redox properties of GbpA-type LPMOs. Our data show that the redox properties of *Vc*GbpA and *Vh*GbpA are very similar. Moreover, the reduction and reoxidation rates obtained for these to GbpA’s are similar to those obtained for the well-characterized bacterial chitin-active LPMO *Sm*AA10A (or CBP21) at pH 7.0 (60). Considering the obtained second-order rate constants for reduction by AscA, even the lowest AscA concentration employed in this study (25 μM) would yield a rate of reduction of 3 – 4 s^-1^, which is similar to the *k*_cat_ of 6.7 s^-1^ determined for the peroxygenase reaction catalyzed by *Sm*AA10A (34) and orders of magnitude higher than the rate of the oxidase reaction. Thus, it is highly unlikely that reduction is rate-limiting in any of the experiments described here.

For comparison, we also assessed reoxidation by O_2_. As expected, based on the observed rates of the oxidase reaction (0.15 min^-1^ to 0.66 min^-1^; see above) and earlier studies (e.g. (60)), reoxidation by O_2_ was found to be exceedingly slow (**Fig. S5**) ranging from 0.05 min^-1^ to 0.07 min^-1^ at ambient conditions (approximately 250 μM O_2_), which, assuming linearity, would translate to second-order rate constants on the order of 3.3 – 4.6 M^-1^⋅s^-1^, i.e., some 1000-fold lower compared to the reoxidation rates obtained with H_2_O_2_. Of note, rate constants for reoxidation by O_2_ are orders of magnitude higher, often amounting to 10^5^ to 10^7^ M^-1^⋅s^-1^, for binuclear copper enzymes such as dopamine monooxygenase (67) and galactose oxidase (68), as well as for certain multicopper laccases (69).

All in all, these data show that *Vc*GbpA and *Vh*GbpA are very similar enzymes and that the copper reactivity of their LPMO domain is identical to that of other chitin-active bacterial LPMOs. The data further suggest that truncation of the CBM hardly affects reactivity, whereas truncation of all three additional domains does seem to modulate the reactivity of the LPMO domain to some extent. This is discussed further below.

### 3.2 Binding to chitin

Previous studies have shown that *Vc*GbpA binds *N*-acetylglucosamine (GlcNAc) containing sugars (11) including different forms of chitin (46). To investigate whether this extends to *Vh*GbpA and to assess the possible impact of the different domains, we studied the binding of the various *Vc*GbpA and *Vh*GbpA variants to chitin in the absence and presence of ascorbate. From earlier studies, it is known that reduction of the LPMO promotes substrate binding (70). Binding to α-chitin was weak in all cases, even in the presence of ascorbate, preventing reliable comparison of the variants (results not shown). In contrast, the binding studies with β-chitin revealed clear binding, and, moreover revealed some remarkable differences between the enzyme variants (**Fig. 2**). As expected, the presence of reductant promoted the binding of *Vh*GbpA^FL^ and *Vh*GbpA^CD^, but this effect was absent for *Vh*GbpA^𝛥CBM73^ (**Fig. 2A, B**). The *Vh*GbpA^𝛥CBM73^ variant bound weakly to the substrate, regardless of the presence of a reductant. Analysis of binding to β-chitin was also analyzed over time for variants of both *Vh*GbpA and *Vc*GbpA. The binding curves (**Fig. 2C, D**) show rapid binding within 1 minute, which was the shortest possible sampling interval, and confirmed the remarkable observation described above: for both *Vh*GbpA and *Vc*GbpA, the CBM73-truncated variant showed weaker binding to chitin, than the catalytic domain only (**Fig. 2C, D**).

**Figure 2.**
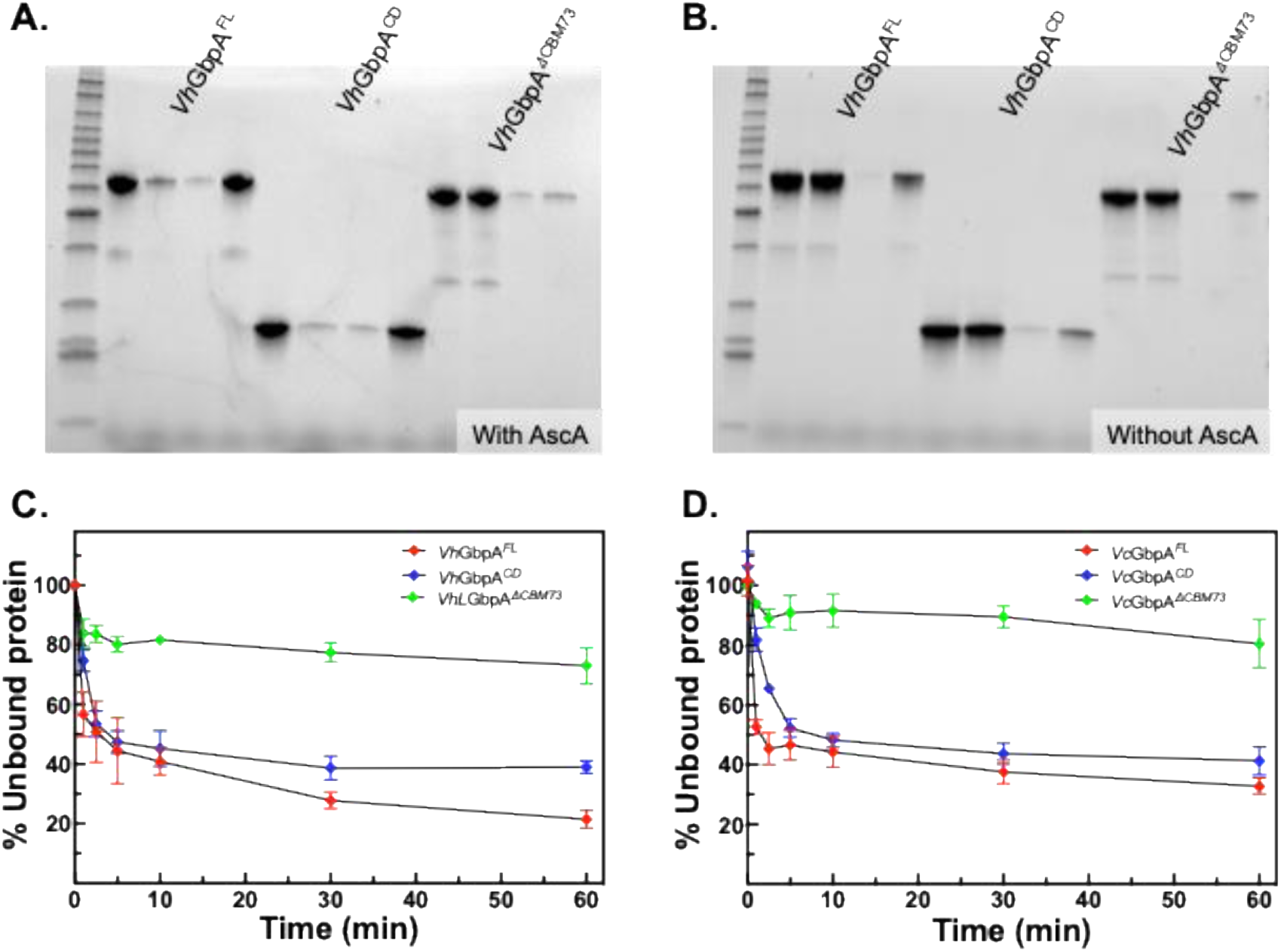
Binding of *Vh*GbpA and *Vc*GbpA variants to β-chitin. (**A, B**) Cu(II)-loaded *Vh*GbpA variants (10 μM for *Vh*GbpA and *Vh*GbpA^𝛥CBM73^, 20 μM for *Vh*GbpA^CD^) were incubated with 10 g.L^-1^ β-chitin in the presence (**A**) or absence (**B**) of 1 mM ascorbic acid in 20 mM Tris-HCl, pH 7.5, at 22°C, 1000 rpm for one hour. Subsequently, the chitin bound enzyme was separated from the unbound fraction by centrifugation. The chitin fraction was washed twice with buffer and then resolubilized in an equal volume of SDS-PAGE sample buffer. The protein content in each fraction was qualitatively analyzed by SDS-PAGE. For each binding reaction, four samples are shown: C., supernatant of a reaction without chitin; S., supernatant of a reaction with chitin; W., supernatant from the second washing step; P., protein in the washed chitin pellet (this protein was solubilized by incubating the pellet in SDS-PAGE sample buffer at 95°C). (**C, D**) Binding of *Vh*GbpA and *Vc*GbpA variants to β-chitin over time in the presence of ascorbic acid. The reactions were performed in 20 mM Tris-HCl, pH 7.5, at 22°C, 1,000 rpm using 2 μM enzyme, 10 g.L^-1^ β-chitin. In this experiment, the non-bound enzyme was separated from the insoluble chitin fraction by filtration using a vacuum manifold and quantified in 96-well microtiter plates using a modified Bradford assay (62).

These results are unexpected for several reasons. Firstly, the fact that the full length enzyme and the catalytic domain only bind approximately equally strongly to chitin is unexpected, considering that the latter lacks the CBM73 with affinity for chitin (54) and considering previous studies showing that chitin-binding CBMs enhance substrate binding by chitin-active LPMOs (42). Secondly, and most remarkably, the removal of the CBM73 somehow prevents the binding of the catalytic domain to chitin. This may be taken to suggest that in the CBM73-truncated variant, the two internal domains, GbpA2 and GbpA3, somehow interact with the substrate-binding surface of the catalytic domain, as discussed in detail below. It is important to note that these effects are much clearer when the catalytic domain is reduced (compare **Fig 2A with 2B)**, i.e., when it is catalytically competent and vulnerable to off-pathway reactions.

### 3.3 Activity and stability during chitin degradation

Substrate-binding affinities of LPMOs can also be assessed by studying enzyme activity and stability in turnover conditions using varying substrate concentrations. Weak binding of the LPMO and/or low substrate concentrations will promote off-pathway reactions, which will lead to enzyme inactivation. In other words, an LPMO that binds its substrate weakly will perform less well at lower substrate concentrations compared to an LPMO that binds the substrate more strongly. It is important to note that in reductant-driven reactions, such as those discussed here, the rate of the reaction primarily depends on the rate of *in situ* generation of H_2_O_2_. Thus, the rate will not necessarily depend on the substrate concentration, even if this concentration is not saturating. Furthermore, inactivation of the LPMO will only become noticeable when the remaining fraction of active enzymes becomes too small to consume the *in situ* generated H_2_O_2_ productively.

In accordance with these latter considerations, an extensive study with the *Vh*GbpA variants using five different chitin concentrations showed that the initial rates of the chitin degradation reactions were similar for all three variants with only a modest dependency on the chitin concentration (**Fig. 3**). However, the progress curves showed very different shapes. The full-length enzyme showed linear progress curves for the full 24 h of the reaction at all substrate concentrations, except the lowest (2.5 g.L^-1^) (**Fig. 3A**). The catalytic domain alone only showed a linear progress curve at the highest tested chitin concentration (50 g/L) and showed increasingly fast deactivation as the substrate concentration became lower (**Fig. 3B**). The CBM73-truncated variant showed clear signs of early inactivation at all tested substrate concentrations (**Fig. 3C**). Due to fast inactivation, product levels in reactions with this latter enzyme remained low, at all tested substrate concentrations

**Figure 3.**
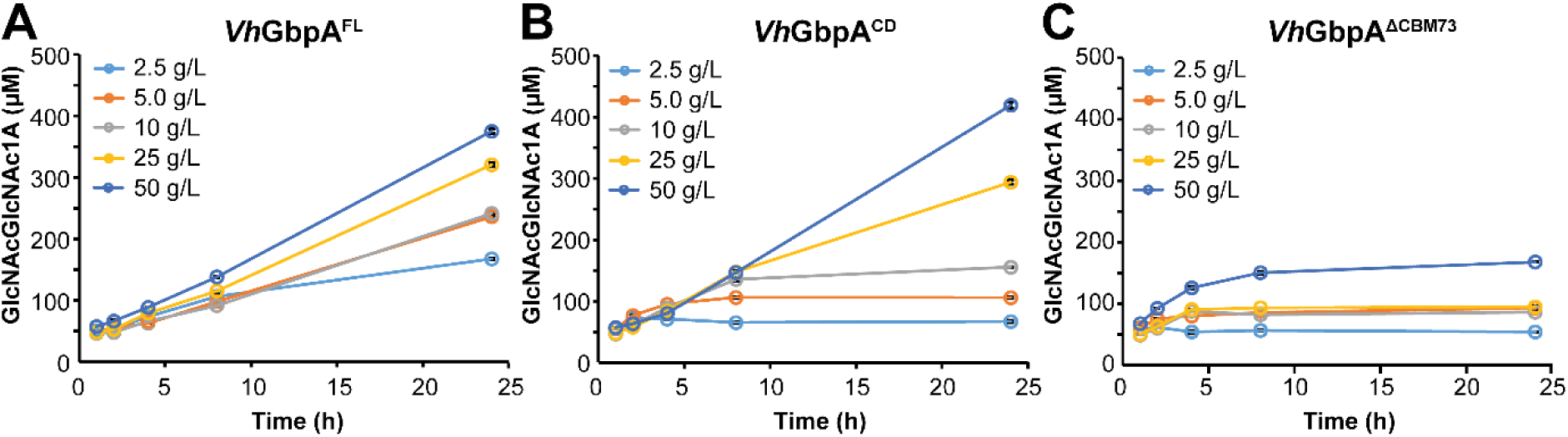
Reductant-driven chitin degradation by variants of *Vh*GbpA at varying substrate concentrations. The graphs show time courses for the formation of soluble oxidized products in reactions containing 0.5 μM *Vh*GbpA^FL^ (**A**) or *Vh*GbpA^CD^ (**B**) or *Vh*GbpA^𝛥CBM73^ (**C**) with β-chitin (2.5-50 g.L^-1^) in 20 mM Tris-HCl buffer, pH 7.5, fueled by 1 mM AscA. Through incubation with *Sm*CHB, soluble products were converted to a mixture of GlcNAc and chitobionic acid (GlcNAcGlcNAc1A) and the latter was quantified to determine the concentration of oxidized products. Each experiment was performed in triplicates; standard deviations were small and are hidden by the symbols.

The observed difference between *Vh*GbpA, with its CBM73, and *Vh*GbpA^CD^ is in line with the results of other studies on the effect of removing substrate-binding domains from two domain LPMOs (38, 43). Substrate binding through the CBM enhances the proximity of the catalytic domain to the substrate, increasing the chance that available H_2_O_2_ is used productively to oxidize the substrate rather than in a futile turnover reaction that may damage the enzyme. In light of existing literature data, the observed difference in activity and stability during chitin degradation between *Vh*GbpA^FL^ and *Vh*GbpA^CD^ is exactly what one would expect upon removal of the CBM.

The results obtained with *Vh*GbpA^𝛥CBM73^ are puzzling but not surprising considering the binding studies described above. If this enzyme variant does not bind well to the substrate, as the binding studies suggest, and if this variant can still react with small molecules in off-pathway reactions, as the assessment of its redox properties suggests, one would expect the enzyme to rapidly inactivate in turnover conditions.

Another way to assess the ability of an LPMO to productively use H_2_O_2_ in a reaction with substrate is to look at progress curves for substrate turnover in reactions with varying amounts of exogenously added H_2_O_2_ at the start of the reaction. The higher the initial H_2_O_2_ concentration and the lower the ability of the LPMO domain to interact productively with the substrate, the faster enzyme inactivation will be observed. Progress curves obtained with different H_2_O_2_ concentrations (**Fig. 4**) show fast turnover of the substrate, in accord with the peroxygenase nature of these enzymes (note the minute time scale in **Fig. 4** compared to the hour time scale in **Fig. 3**). In terms of tolerance to H_2_O_2_ and enzyme inactivation, the progress curves for both *Vh*GbpA (**Fig. 4**) and *Vc*GbpA (**Fig. S6**) show trends that align well with the trends observed for the reductant-driven reactions.

**Figure 4.**
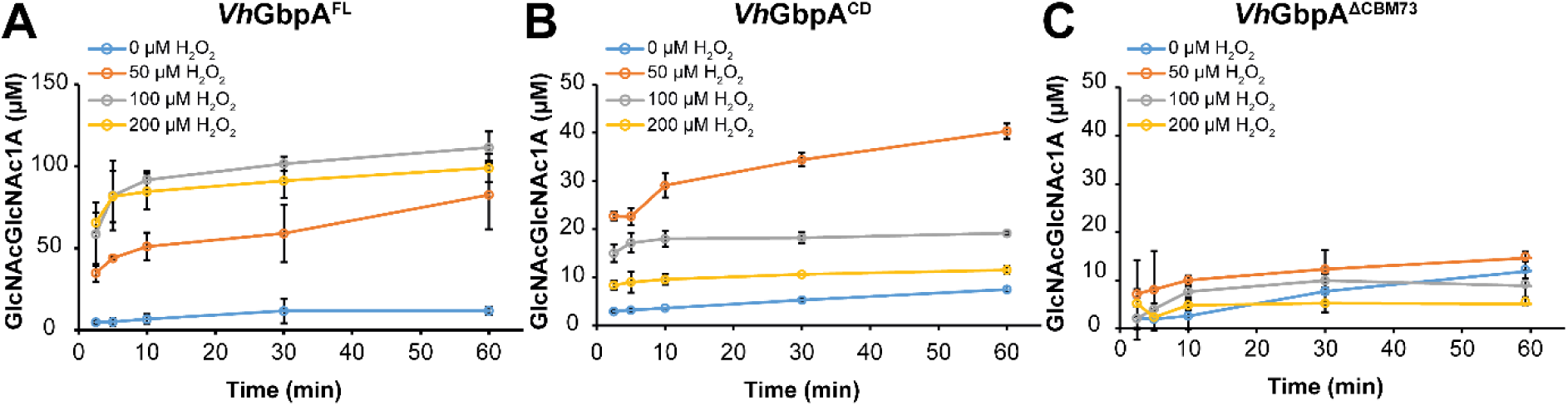
H_2_O_2_-driven degradation of chitin by *Vh*GbpA variants. The graphs show time courses for the formation of soluble oxidized products in reactions containing 0.5 µM *Vh*GbpA^FL^ (**A**), *Vh*GbpA^CD^ (**B**) or *Vh*GbpA^𝛥CBM73^(**C**), 10 g.L^-1^ β-chitin in 20 mM Tris-HCl, pH 7.5. After pre-incubating these mixtures at 30°C with shaking at 1000 rpm for 30 mins, reactions were initiated by sequentially adding H_2_O_2_ (to the indicated final concentrations) and, lastly, 0.1mM AscA to start the reaction. Soluble oxidized products were quantified as described in the legend of **Fig. 3**. All reactions were performed in triplicates and the error bars show ± S.D. (n = 3). Note that product formation in the reaction with 0 μM H_2_O_2_ reflects reductant-driven LPMO activity; in this reaction, H_2_O_2_ is generated slowly, *in situ*. Identical experiments with the *Vc*GbpA variants gave similar results and are shown in **Fig. S6**.

Firstly, **Fig. 4** shows that the full-length enzyme is more capable of productively oxidizing chitin than the catalytic domain only. For the full-length enzyme, the reactions with 100 and 200 µM H_2_O_2_ both yielded about 100 µM of product, showing that notable inactivation only occurs at 200 µM H_2_O_2_. For the catalytic domain, product formation at 100 and 200 µM H_2_O_2_ amounted to approximately 20 and 10 µM, respectively, which is lower than the amount of product obtained in the reaction with 50 µM H_2_O_2_. This confirms the notion that the catalytic domain only, lacking the CBM, is less capable of productively using H_2_O_2_ and more prone to damaging off pathway reactions. Secondly, **Fig. 4C** shows that the CBM-truncated variant is rapidly inactivated at all H_2_O_2_ concentrations, confirming that the productive interaction of this enzyme with the substrate is severely hampered.

### 3.4 Stability of the reduced enzyme in the absence of substrate

Given that reduced LPMOs are prone to oxidative self-inactivation and that the *Vh*GbpA variants show clear differences in catalytic performance, we assessed their stability in the absence of substrate using an ascorbic acid depletion assay (**Fig. 5**). In this experiment, AscA depletion serves as a proxy for copper release following enzyme damage and inactivation, processes that are triggered under reductive, substrate-free conditions (38, 45). In such conditions, the reduced LPMO will react with available H_2_O_2_ in a damaging off-pathway reaction and copper released from damaged active sites will promote oxidation (depletion) of AscA. Thus, this experiment may show to what extent the GbpA variants are protected from turning over H_2_O_2_ when there is no substrate present. The full-length enzyme showed the slowest consumption of AscA suggesting that it is most stable and least likely to turn over H_2_O_2_ in the absence of substrate. The CBM73-truncated variant was somewhat less stable than the full-length enzyme, whereas, importantly, the catalytic domain, clearly was the least stable of the three GbpA variants. This shows that while the catalytic domain alone binds chitin and retains high activity, it is also more vulnerable to oxidative inactivation without the stabilizing contribution of the accessory domains GbpA2 and GbpA3. A plausible explanation for this observation, aligning with similar indications described above, would be that the GbpA2 and GbpA3 domains interact with the reduced copper site on the LPMO domain, thus reducing its tendency to turnover H_2_O_2_ in the absence of substrate.

**Figure 5.**
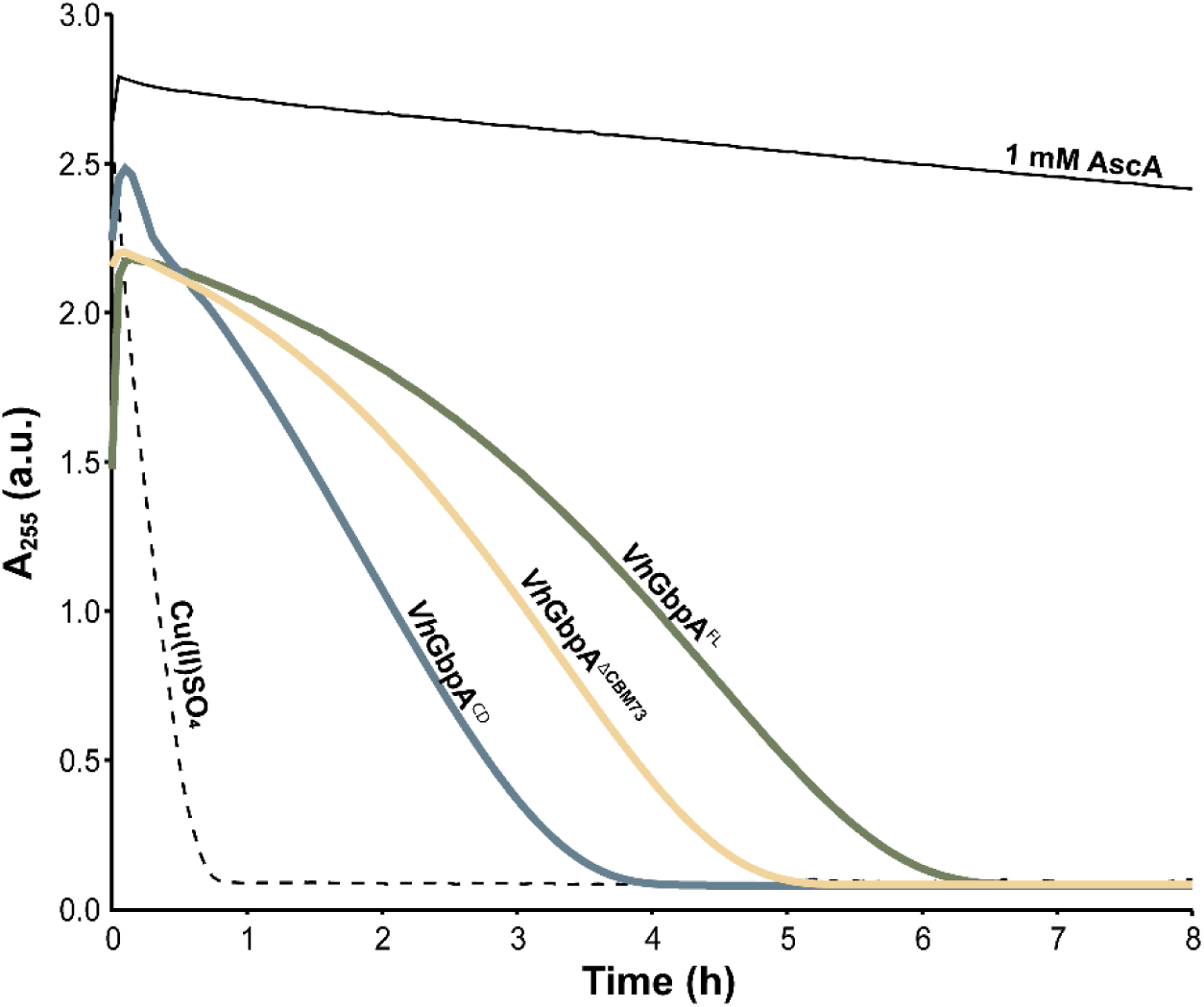
Ascorbic acid depletion in substrate-free reactions with *Vh*GbpA variants. Reactions containing 0.5 µM of each *Vh*GbpA variant were performed in 20 mM Tris-HCl buffer (pH 7.5) with 1 mM ascorbic acid (AscA). A control with 0.5 µM Cu(II)SO_4_ was included to monitor the effect of free copper, and an additional control lacking enzyme (1 mM AscA in buffer) was used to assess AscA stability over time. All reactions were carried out in technical triplicates (n = 3), but for clarity only one representative curve per variant is shown, as replicates showed negligible variation. The mixtures were incubated at 30 °C for up to 8 hours, and absorbance at 255 nm was recorded every 3 minutes. Gentle mixing was achieved through plate movement during measurements.

### 3.5 An interaction between GbpA3 and the catalytic domain?

The crystal structures of CBM73 truncated *Vc*GbpA (46) and of full-length *Vh*GbpA (51) do not point at specific interactions between the GbpA2 or GbpA3 domains and the substrate-binding surface and catalytic centre of the LPMO domain. SAXS studies of both enzymes suggest that they have an elongated form in solution (46, 51). Nevertheless, the present data clearly indicate that when the CBM73 is truncated, the presence of the GbpA2 and GbpA3 domains hampers binding to chitin Importantly, crystallization and SAXS studies of the GbpAs have been done using enzymes in their Cu(II) state, whereas the remarkable findings described above refer to GbpAs in their Cu(I) state. The AscA depletion assay, where the LPMO is in the Cu(I) state also suggests an interaction between the domains.

To gain more insight into these matters, we predicted the structures of both *apo*- and copper-loaded GbpAs using the AlphaFold3 online server (56). Most interestingly, the models predict that GbpA3 folds back onto the substrate-biding surface of the LPMO domain and that two surfaces exposed histidines (His335 and His366 in *Vh*GbpA and His334 and His365 in *Vc*GbpA) in GbpA3 can interact with the copper in the histidine brace (**Fig. 6 & Fig. S7**). This interaction was only predicted when copper was included, and the absence or presence of the CBM73 did not change the outcome of these predictions. This suggests a potential regulatory role for the GbpA3 domain in modulating access to the copper centre of the LPMO domain, which aligns well with the experimental data described above.

**Figure 6.**
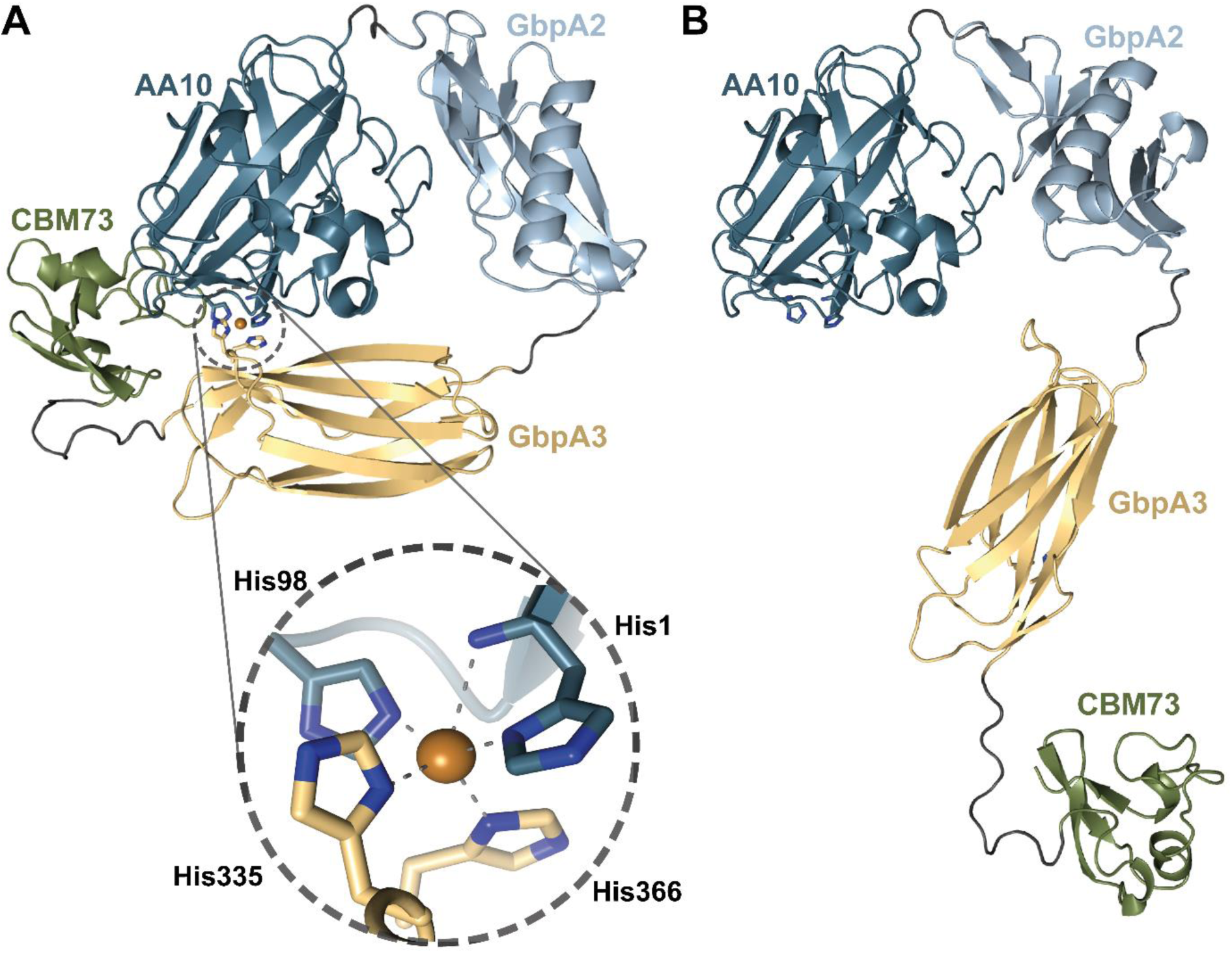
Cartoon representations of the structures of *holo* and *apo Vh*GbpA predicted by AlphaFold3. Panel A shows the copper-loaded (*holo*) *Vh*GbpA structure, where the GbpA3 domain folds over the AA10 catalytic site. In this conformation, two histidine residues from GbpA3 (His335 and His366) are positioned to coordinate the copper ion in the active site together with the histidine-brace (His1 & His98). Panel B displays the *apo*-*Vh*GbpA structure, which adopts a more elongated conformation. A similar domain rearrangement upon copper binding is observed when comparing AlphaFold3 predictions for the structure of *apo* and *holo-Vc*GbpA (**Fig. S7**).

An analysis of 165 GbpA-like protein sequences (InterPro IPR020879), each containing all four domains, revealed that the two histidines predicted to interact with the active site copper in *Vh*GbpA3 and *Vc*GbpA are strictly conserved across all sequences (**Fig. S8**). Interestingly, for these 165 proteins, sequence conservation is markedly higher in the AA10 (LPMO) domain than in the GbpA3 domain (**Fig. S9**). While the AA10 domain shows full conservation for 60 out of 182 positions, only 7 out of 89 residues in the GbpA3 domain are fully conserved and two of these are the histidines that are putatively interacting with the catalytic copper. An additional search in InterPro for proteins containing the GbpA3 domain led to the discovery of 28 multimodular GH18 chitinases harboring a GbpA3 domain, including a GH18 (UniProt ID A7MTY3) from *V. campbellii* ATCC BAA-1116, which is the source of the *Vh*GbpA studied here. Analysis of these non-LPMO GbpA3 domains revealed that the histidines conserved in GbpA-like LPMO proteins are absent. In fact, none of these two histidines were present in any of the GbpA3 domains of these GH18-containing proteins. The positions corresponding to His335 and His366 in *Vh*GbpA3 are instead occupied by a serine or an asparagine (His335 position) and a glutamine or a serine (His366 position), respectively. Thus, the presence of these two histidines in GbpA3 domains is correlated with the GbpA3 domain being part of an LPMO.

A further analysis of the interaction between the AA10 domain and the GbpA3 domain revealed other highly conserved residues on the AA10 domain and the GbpA3 domain that could contribute to the interaction between these domains (**Fig. S10**). For example, the side chain of Glu39 located on the substrate binding surface of the LPMO domain can form a hydrogen bond with the side chain of Gln383 on the GbpA3 domain, whilst the main chain carbonyl of Gln383 can form a hydrogen bond with the side chain of Arg20. All these residues are fully conserved, except the arginine at position 20, which is an exception; the other 164 GbpAs have a lysine, which can make a similar hydrogen bond (**Fig. S9).** Finally, fully conserved Glu44 and Thr166 on the AA10 domain can form hydrogen bonds to the imidazole side chain of His335 (**Fig. S10**).

The combination of these structural predictions with the observation that chitin binding is only hampered by the GbpA3 domain in the reduced LPMO and, most remarkably, only in the absence of the CBM73, leads to a hypothesis for how the reactivity of the catalytically competent copper site may be modulated by the presence of substrate (**Fig. 7**). One could envisage that in the absence of substrate the reduced copper in the catalytic LPMO domain is protected from engaging in off-pathway reactions by binding of GbpA3, as supported by, for example, the AscA depletion assay. Binding of the CBM73 to substrate could lead to elongation of the protein, pulling GbpA3 away from the LPMO domain, enabling the latter to interact with the substrate. In the CBM73-truncated variants, this “pulling effect” would not occur, explaining why these variants interact badly with chitin.

**Figure 7.**
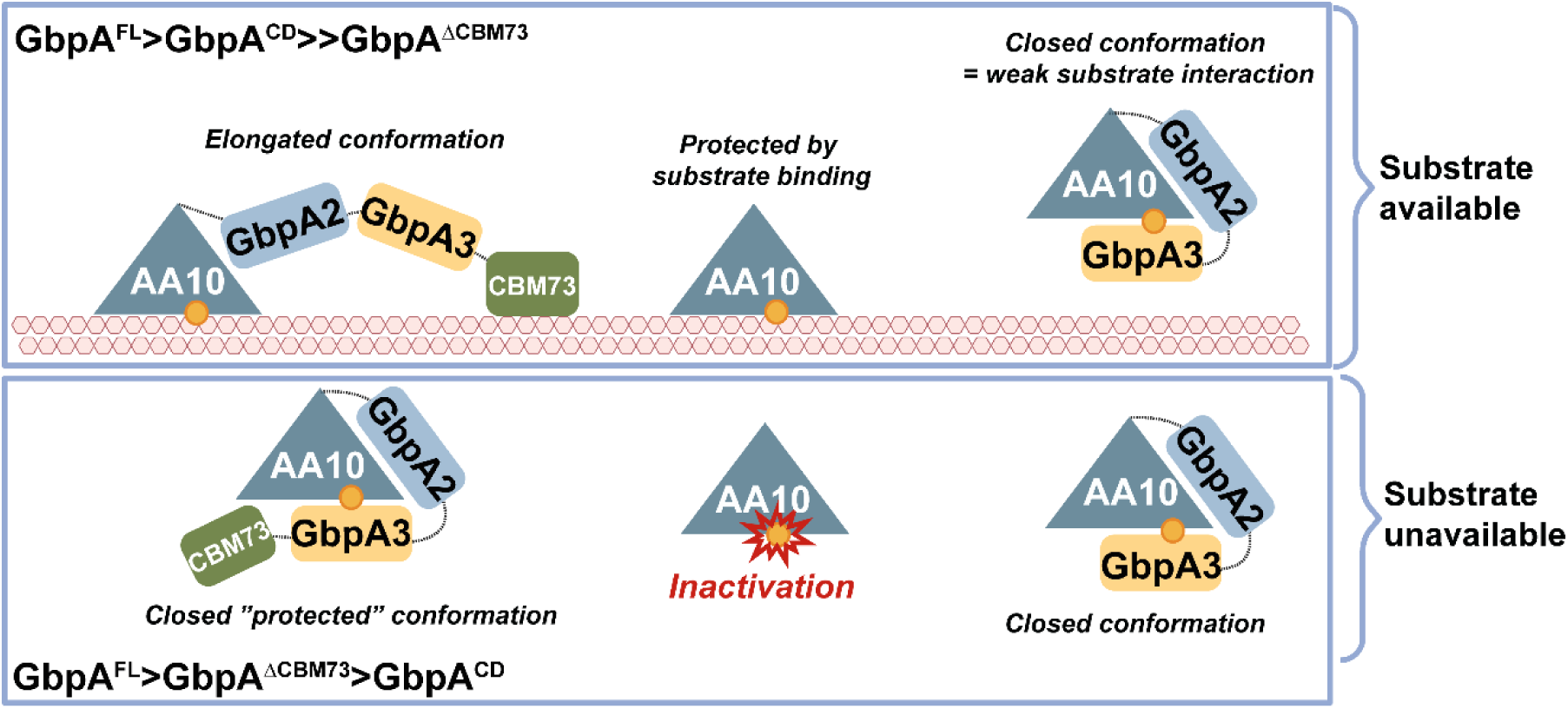
Proposed domain arrangement of GbpA in the presence and absence of chitin. When chitin is present, the catalytic AA10 domain and the CBM73 domain are predicted to bind the chitin surface, resulting in an elongated conformation that displaces the GbpA3 domain from the LPMO active site, thereby facilitating productive interactions between the LPMO and the substrate. In the absence of chitin, the model predicts that GbpA3 folds back over the substrate-binding surface of the LPMO. In this conformation, two surface-exposed histidines (His335 and His366 in *Vh*GbpA; His334 and His365 in *Vc*GbpA, see **Fig. 6** and **S7**) are positioned to interact with the catalytic copper ion within the histidine brace. In CBM73-truncated variants, the absence of both the CBM domain and substrate binding likely prevents the protein from adopting the elongated, active conformation, which explains their reduced ability to act on chitin and increase susceptibility to inactivation (relative to the other two enzyme variants in the presence of substrate). In the absence of substrate, the LPMO domain alone is more rapidly inactivated (indicated by the red symbol) than the CBM73-truncated variant because of the protective function of the GbpA3 domain in the latter. The order of the stabilities of the GbpA variants in the presence (upper panel) or absence (lower panel) of substrate is indicated in the Figure.

Interestingly, a preprint by Sørensen *et al.* (71) addresses the conformation of *Vc*GbpA when bound to chitin. Using negative-stain electron microscopy, Sørensen *et al.* conclude that *Vc*GbpA adopts an extended conformation when bound to chitin, well in line with the model proposed in **Fig. 7**.

The interaction of the GbpA3 domain with the LPMO domain is strongly supported by the impact of GbpA3 on binding and degradation of chitin and by most of the additional functional analyses that assess copper reactivity in the absence of substrate. The observed higher oxidase activity for the catalytic domain alone compared to the two variants carrying the GbpA3 domain aligns very well with the hypothesis that GbpA3 affects the reactivity of the LPMO domain. However, single-turnover experiments for reduction by AscA and reoxidation by H₂O₂ did not show similarly large effects. It is conceivable that this reflects a limitation of the “single-turnover” set-up, although the exact nature of such an effect remains elusive. While differences observed in the reduction and reoxidation rates were small (**Table 1**), these differences do in fact align with the rest of the data and our hypotheses: Firstly, GbpA variants with the GbpA3 domain show the faster reduction rates, which may be due to a favorable interaction between the reduced LPMO domain and the GbpA3 domain that promotes reduction. Secondly, the LPMO domain alone shows the faster reoxidation rate, compared to the GbpA3 containing enzyme variants, which aligns with the notion that the LPMO is less protected in the absence of the GbpA3 domain. Finally, the AscA depletion assays support the notion that, in absence of substrate, the GbpA3 domain protects the copper from reacting with H_2_O_2_. Of note, this latter experiment may seem to suggest that the presence of the CBM73 somehow affects the LPMO-GbpA3 interaction, since *Vh*GbpA^FL^ and *Vh*GbpA^𝛥CBM73^, both with the GbpA3 domain, show different depletion curves.

## 4. Concluding remarks

This study provides an in-depth functional characterization of an essential class of multi-domain LPMOs with roles in bacterial virulence. While the LPMO domain of these GbpA proteins turns out to behave similarly to single-domain chitin-active LPMOs, the multi-domain structure of these proteins sets them apart from LPMOs that seem to have evolved to degrade chitin, such as *Sm*AA10A from *S. marcescens* (15, 72). It is worth noting that multi-domain LPMOs associated with virulence occur in ecosystems where degradation of chitin does not seem to occur (55), and it may be questioned whether the generally observed activity on chitin for these proteins reflects their natural substrate preferences or whether these enzymes may have other, hitherto unknown substrates.

Our functional characterization, combined with structure predictions, clearly shows that the GbpA3 domain binds to the reduced LPMO domain. This provides a glimpse of the function of this domain, which we believe is linked to regulating the access to and reactivity of the copper site, as illustrated in **Fig.6**. This unique arrangement provides a novel concept in the LPMO field and may eventually contribute to understanding how these multi-domain LPMO-containing virulence factors operate. We note that GbpA3-like proteins are widely distributed across bacterial species, with conserved sequence motifs suggesting functional relevance across diverse ecosystems.

It remains to be seen how this interaction mechanism plays out in natural environments. Does this protein primarily act on chitin, or are other substrates involved? Investigating CBM73 specificity and conducting *in vivo* experiments with truncation variants will be crucial to resolving these questions. Furthermore, investigating if and how GbpA2 and GbpA3 interact with chitin surfaces, or potentially other biomolecular structures, could provide broader insights into their roles in substrate recognition, enzymatic efficiency, and into their ecological relevance.

To better understand the interaction between GbpA3 and the LPMO domain, as well as the functional implications of this interaction, additional work is needed. This could include affinity studies using isolated domains, site-directed mutagenesis, and investigations into conformational changes upon substrate binding. It would also be of significant interest to study how this newly discovered domain-domain interaction affects GbpA functionality *in vivo*. Despite being around for two decades now, many questions remain regarding the *in vivo* functions of these intriguing virulence factors. The present results suggest that the unique domain arrangement in GbpA-like proteins not only relates to substrate binding but also to protecting the enzyme from damaging off-pathway reactions.

## 5. Acknowledgements & Funding

This work was sponsored by the Vidyasirimedhi Institute of Science and Technology (VISTEC) through a full Ph.D. scholarship to YZ. WS received supportive funding by the National Research Council of Thailand (NRCT) through Mid-Career Grant No. N42A660311 (B23WIS-NRC010). VGHE, EK, and ZF were supported by an ERC-Synergy grant (856446). YZ acknowledges the kind continuous assistance from the PEP lab members during his one-year research visit at KBM, NMBU.

## References

1. Blackwell, J. (1969) Structure of β-chitin or parallel chain systems of poly-β-(1-4)-*N*-acetyl-D-glucosamine Biopolymers 7, 281–298

2. Rinaudo, M. (2006) Chitin and chitosan: Properties and applications Progress in Polymer Science 31, 603–632

3. Kurita, K. (2006) Chitin and chitosan: functional biopolymers from marine crustaceans Mar Biotechnol (NY) 8, 203–226

4. Lipke, P. N., and Ovalle, R. (1998) Cell wall architecture in yeast: new structure and new challenges J Bacteriol 180, 3735–3740

5. Bowman, S. M., and Free, S. J. (2006) The structure and synthesis of the fungal cell wall Bioessays 28, 799–808

6. Amiri, H., Aghbashlo, M., Sharma, M., Gaffey, J., Manning, L., Moosavi Basri, S. M. et al. (2022) Chitin and chitosan derived from crustacean waste valorization streams can support food systems and the UN Sustainable Development Goals Nat Food 3, 822–828

7. Souza, C. P., Almeida, B. C., Colwell, R. R., and Rivera, I. N. (2011) The importance of chitin in the marine environment Mar Biotechnol (NY) 13, 823–830

8. Devlin, J. R., and Behnsen, J. (2023) Bacterial chitinases and their role in human infection Infect Immun 91, e0054922

9. Jeong, G. J., Khan, F., Tabassum, N., and Kim, Y. M. (2023) Chitinases as key virulence factors in microbial pathogens: Understanding their role and potential as therapeutic targets Int J Biol Macromol 249, 126021

10. Stauder, M., Huq, A., Pezzati, E., Grim, C. J., Ramoino, P., Pane, L. et al. (2012) Role of GbpA protein, an important virulence-related colonization factor, for Vibrio cholerae’s survival in the aquatic environment Environ Microbiol Rep 4, 439–445

11. Kirn, T. J., Jude, B. A., and Taylor, R. K. (2005) A colonization factor links Vibrio cholerae environmental survival and human infection Nature 438, 863–866

12. Askarian, F., Uchiyama, S., Masson, H., Sorensen, H. V., Golten, O., Bunaes, A. C., et al. (2021) The lytic polysaccharide monooxygenase CbpD promotes Pseudomonas aeruginosa virulence in systemic infection Nat Commun 12, 1230

13. Askarian, F., Tsai, C. M., Cordara, G., Zurich, R. H., Bjanes, E., Golten, O., et al. (2023) Immunization with lytic polysaccharide monooxygenase CbpD induces protective immunity against Pseudomonas aeruginosa pneumonia Proc Natl Acad Sci U S A 120, e2301538120

14. Vaaje-Kolstad, G., Horn, S. J., van Aalten, D. M., Synstad, B., and Eijsink, V. G. (2005) The non-catalytic chitin-binding protein CBP21 from Serratia marcescens is essential for chitin degradation J Biol Chem 280, 28492–28497

15. Vaaje-Kolstad, G., Westereng, B., Horn, S. J., Liu, Z., Zhai, H., Sorlie, M. et al. (2010) An oxidative enzyme boosting the enzymatic conversion of recalcitrant polysaccharides Science 330, 219–222

16. Vaaje-Kolstad, G., Horn, S. J., Sorlie, M., and Eijsink, V. G. (2013) The chitinolytic machinery of Serratia marcescens - a model system for enzymatic degradation of recalcitrant polysaccharides FEBS J 280, 3028–3049

17. Charoenpol, A., Crespy, D., Schulte, A., and Suginta, W. (2023) Marine chitin upcycling with immobilized chitinolytic enzymes: current state and prospects Green Chemistry 25, 467–489

18. Vandhana, T. M., Reyre, J. L., Sushmaa, D., Berrin, J. G., Bissaro, B., and Madhuprakash, J. (2022) On the expansion of biological functions of lytic polysaccharide monooxygenases New Phytol 233, 2380–2396

19. Quinlan, R. J., Sweeney, M. D., Lo Leggio, L., Otten, H., Poulsen, J. C., Johansen, K. S., et al. (2011) Insights into the oxidative degradation of cellulose by a copper metalloenzyme that exploits biomass components Proc Natl Acad Sci U S A 108, 15079–15084

20. Phillips, C. M., Beeson, W. T., Cate, J. H., and Marletta, M. A. (2011) Cellobiose dehydrogenase and a copper-dependent polysaccharide monooxygenase potentiate cellulose degradation by Neurospora crassa ACS Chem Biol 6, 1399–1406

21. Bissaro, B., Rohr, A. K., Muller, G., Chylenski, P., Skaugen, M., Forsberg, Z. et al. (2017) Oxidative cleavage of polysaccharides by monocopper enzymes depends on H_2_O_2_ Nat Chem Biol 13, 1123–1128

22. Munzone, A., Eijsink, V. G. H., Berrin, J. G., and Bissaro, B. (2024) Expanding the catalytic landscape of metalloenzymes with lytic polysaccharide monooxygenases Nat Rev Chem 8, 106–119

23. Hagemann, M. M., and Hedegard, E. D. (2023) Molecular mechanism of substrate oxidation in lytic polysaccharide monooxygenases: Insight from theoretical investigations Chemistry 29, e202202379

24. Hagemann, M. M., Wieduwilt, E. K., Ryde, U., and Hedegard, E. D. (2024) Investigating the substrate oxidation mechanism in lytic polysaccharide monooxygenase: H_2_O_2_- versus O_2_- activation Inorg Chem 63, 21929–21940

25. Lim, H., Brueggemeyer, M. T., Transue, W. J., Meier, K. K., Jones, S. M., Kroll, T., et al. (2023) Kβ X-ray emission spectroscopy of Cu(I)-lytic polysaccharide monooxygenase: Direct observation of the frontier molecular orbital for H_2_O_2_ activation J Am Chem Soc 145, 16015–16025

26. Wang, B., Wang, Z., Davies, G. J., Walton, P. H., and Rovira, C. (2020) Activation of O_2_ and H_2_O_2_ by lytic polysaccharide monooxygenases ACS Catalysis 10, 12760–12769

27. Muller, G., Varnai, A., Johansen, K. S., Eijsink, V. G., and Horn, S. J. (2015) Harnessing the potential of LPMO-containing cellulase cocktails poses new demands on processing conditions Biotechnol Biofuels 8, 187

28. Guo, X., An, Y., Liu, F., Lu, F., and Wang, B. (2022) Lytic polysaccharide monooxygenase - A new driving force for lignocellulosic biomass degradation Bioresour Technol 362, 127803

29. Hemsworth, G. R., Johnston, E. M., Davies, G. J., and Walton, P. H. (2015) Lytic polysaccharide monooxygenases in biomass conversion Trends Biotechnol 33, 747–761

30. Span, E. A., Suess, D. L. M., Deller, M. C., Britt, R. D., and Marletta, M. A. (2017) The role of the secondary coordination sphere in a fungal polysaccharide monooxygenase ACS Chem Biol 12, 1095–1103

31. Hall, K. R., Joseph, C., Ayuso-Fernandez, I., Tamhankar, A., Rieder, L., Skaali, R. et al. (2023) A conserved second sphere residue tunes copper site reactivity in lytic polysaccharide monooxygenases J Am Chem Soc 145, 18888–18903

32. Jones, S. M., Transue, W. J., Meier, K. K., Kelemen, B., and Solomon, E. I. (2020) Kinetic analysis of amino acid radicals formed in H_2_O_2_-driven Cu(I) LPMO reoxidation implicates dominant homolytic reactivity Proc Natl Acad Sci U S A 117, 11916–11922

33. Rieder, L., Petrovic, D., Valjamae, P., Eijsink, V. G. H., and Sorlie, M. (2021) Kinetic characterization of a putatively chitin-active LPMO reveals a preference for soluble substrates and absence of monooxygenase activity ACS Catal 11, 11685–11695

34. Kuusk, S., Bissaro, B., Kuusk, P., Forsberg, Z., Eijsink, V. G. H., Sorlie, M. et al. (2018) Kinetics of H_2_O_2_-driven degradation of chitin by a bacterial lytic polysaccharide monooxygenase J Biol Chem 293, 523–531

35. Schwaiger, L., Csarman, F., Chang, H., Golten, O., Eijsink, V. G. H., and Ludwig, R. (2024) Electrochemical monitoring of heterogeneous peroxygenase reactions unravels LPMO kinetics ACS Catal 14, 1205–1219

36. Hedison, T. M., Breslmayr, E., Shanmugam, M., Karnpakdee, K., Heyes, D. J., Green, A. P. et al. (2021) Insights into the H_2_O_2_-driven catalytic mechanism of fungal lytic polysaccharide monooxygenases FEBS J 288, 4115–4128

37. Stepnov, A. A., Forsberg, Z., Sorlie, M., Nguyen, G. S., Wentzel, A., Rohr, A. K., et al. (2021) Unraveling the roles of the reductant and free copper ions in LPMO kinetics Biotechnol Biofuels 14, 28

38. Stepnov, A. A., Eijsink, V. G. H., and Forsberg, Z. (2022) Enhanced in situ H_2_O_2_ production explains synergy between an LPMO with a cellulose-binding domain and a single-domain LPMO Sci Rep 12, 6129

39. Golten, O., Ayuso-Fernandez, I., Hall, K. R., Stepnov, A. A., Sorlie, M., Rohr, A. K. et al. (2023) Reductants fuel lytic polysaccharide monooxygenase activity in a pH-dependent manner. FEBS Lett 597, 1363–1374

40. Hemsworth, G. R. (2023) Revisiting the role of electron donors in lytic polysaccharide monooxygenase biochemistry Essays Biochem 67, 585–595

41. Eijsink, V. G. H., Petrovic, D., Forsberg, Z., Mekasha, S., Rohr, A. K., Varnai, A., et al. (2019) On the functional characterization of lytic polysaccharide monooxygenases (LPMOs) Biotechnol Biofuels 12, 58

42. Forsberg, Z., and Courtade, G. (2023) On the impact of carbohydrate-binding modules (CBMs) in lytic polysaccharide monooxygenases (LPMOs) Essays Biochem 10.1042/EBC20220162

43. Gao, W., Li, T., Zhou, H., Ju, J., and Yin, H. (2024) Carbohydrate-binding modules enhance H_2_O_2_ tolerance by promoting lytic polysaccharide monooxygenase active site H_2_O_2_ consumption J Biol Chem 300, 105573

44. Stepnov, A. A., Christensen, I. A., Forsberg, Z., Aachmann, F. L., Courtade, G., and Eijsink, V. G. H. (2022) The impact of reductants on the catalytic efficiency of a lytic polysaccharide monooxygenase and the special role of dehydroascorbic acid FEBS Lett 596, 53–70

45. Østby, H., Tuveng, T. R., Stepnov, A. A., Vaaje-Kolstad, G., Forsberg, Z., and Eijsink, V. G. H. (2023) Impact of copper saturation on lytic polysaccharide monooxygenase performance ACS Sustain Chem Eng 11, 15566–15576

46. Wong, E., Vaaje-Kolstad, G., Ghosh, A., Hurtado-Guerrero, R., Konarev, P. V., Ibrahim, A. F., et al. (2012) The *Vibrio cholerae* colonization factor GbpA possesses a modular structure that governs binding to different host surfaces PLoS Pathog 8, e1002373

47. Forsberg, Z., Tuveng, T. R., and Eijsink, V. G. H. (2024) A modular enzyme with combined hemicellulose-removing and LPMO activity increases cellulose accessibility in softwood FEBS J 10.1111/febs.17250

48. Mekasha, S., Tuveng, T. R., Askarian, F., Choudhary, S., Schmidt-Dannert, C., Niebisch, A. et al. (2020) A trimodular bacterial enzyme combining hydrolytic activity with oxidative glycosidic bond cleavage efficiently degrades chitin J Biol Chem 295, 9134–9146

49. Paspaliari, D. K., Loose, J. S., Larsen, M. H., and Vaaje-Kolstad, G. (2015) Listeria monocytogenes has a functional chitinolytic system and an active lytic polysaccharide monooxygenase FEBS J 282, 921–936

50. Mutahir, Z., Mekasha, S., Loose, J. S. M., Abbas, F., Vaaje-Kolstad, G., Eijsink, V. G. H. et al. (2018) Characterization and synergistic action of a tetra-modular lytic polysaccharide monooxygenase from Bacillus cereus FEBS Lett 592, 2562–2571

51. Zhou, Y., Wannapaiboon, S., Prongjit, M., Pornsuwan, S., Sucharitakul, J., Kamonsutthipaijit, N. et al. (2023) Structural and binding studies of a new chitin-active AA10 lytic polysaccharide monooxygenase from the marine bacterium Vibrio campbellii Acta Crystallogr D Struct Biol 79, 479–497

52. Manjeet, K., Madhuprakash, J., Mormann, M., Moerschbacher, B. M., and Podile, A. R. (2019) A carbohydrate binding module-5 is essential for oxidative cleavage of chitin by a multi-modular lytic polysaccharide monooxygenase from Bacillus thuringiensis serovar kurstaki Int J Biol Macromol 127, 649–656

53. Loose, J. S., Forsberg, Z., Fraaije, M. W., Eijsink, V. G., and Vaaje-Kolstad, G. (2014) A rapid quantitative activity assay shows that the Vibrio cholerae colonization factor GbpA is an active lytic polysaccharide monooxygenase FEBS Lett 588, 3435–3440

54. Forsberg, Z., Nelson, C. E., Dalhus, B., Mekasha, S., Loose, J. S., Crouch, L. I. et al. (2016) Structural and functional analysis of a lytic polysaccharide monooxygenase important for efficient utilization of chitin in Cellvibrio japonicus J Biol Chem 291, 7300-7312

55. Banse, A. V., VanBeuge, S., Smith, T. J., Logan, S. L., and Guillemin, K. (2023) Secreted Aeromonas GlcNAc binding protein GbpA stimulates epithelial cell proliferation in the zebrafish intestine Gut Microbes 15, 2183686

56. Abramson, J., Adler, J., Dunger, J., Evans, R., Green, T., Pritzel, A. et al. (2024) Accurate structure prediction of biomolecular interactions with AlphaFold 3 Nature 630, 493–500

57. Manoil, C., and Beckwith, J. (1986) A genetic approach to analyzing membrane protein topology Science 233, 1403–1408

58. Mekasha, S., Byman, I. R., Lynch, C., Toupalová, H., Anděra, L., Næs, T. et al. (2017) Development of enzyme cocktails for complete saccharification of chitin using mono-component enzymes from Serratia marcescens. Process Biochemistry 56, 132–138

59. Bissaro, B., Forsberg, Z., Ni, Y., Hollmann, F., Vaaje-Kolstad, G., and Eijsink, V. G. H. (2016) Fueling biomass-degrading oxidative enzymes by light-driven water oxidation. Green Chemistry 18, 5357–5366

60. Bissaro, B., Streit, B., Isaksen, I., Eijsink, V. G. H., Beckham, G. T., DuBois, J. L. et al. (2020) Molecular mechanism of the chitinolytic peroxygenase reaction Proc Natl Acad Sci U S A 117, 1504–1513

61. Bradford, M. M. (1976) A rapid and sensitive method for the quantitation of microgram quantities of protein utilizing the principle of protein-dye binding Anal Biochem 72, 248–254

62. Ernst, O., and Zor, T. (2010) Linearization of the bradford protein assay J Vis Exp 10.3791/1918

63. Heuts, D. P., Winter, R. T., Damsma, G. E., Janssen, D. B., and Fraaije, M. W. (2008) The role of double covalent flavin binding in chito-oligosaccharide oxidase from Fusarium graminearum. Biochem J 413, 175–183

64. Kittl, R., Kracher, D., Burgstaller, D., Haltrich, D., and Ludwig, R. (2012) Production of four Neurospora crassa lytic polysaccharide monooxygenases in Pichia pastoris monitored by a fluorimetric assay Biotechnol Biofuels 5, 79

65. Huynh, K., and Partch, C. L. (2015) Analysis of protein stability and ligand interactions by thermal shift assay Curr Protoc Protein Sci 79, 28.29.21–28.29.14

66. Ayuso-Fernandez, I., Emrich-Mills, T. Z., Haak, J., Golten, O., Hall, K. R., Schwaiger, L. et al. (2024) Mutational dissection of a hole hopping route in a lytic polysaccharide monooxygenase (LPMO) Nat Commun 15, 3975

67. Brenner, M. C., and Klinman, J. P. (1989) Correlation of copper valency with product formation in single turnovers of dopamine beta-monooxygenase Biochemistry 28, 4664–4670

68. Whittaker, M. M., Ballou, D. P., and Whittaker, J. W. (1998) Kinetic isotope effects as probes of the mechanism of galactose oxidase Biochemistry 37, 8426–8436

69. Cole, J. L., Ballou, D. P., and Solomon, E. I. (1991) Spectroscopic characterization of the peroxide intermediate in the reduction of dioxygen catalyzed by the multicopper oxidases J Am Chem Soc 113, 8544–8546

70. Kracher, D., Andlar, M., Furtmuller, P. G., and Ludwig, R. (2018) Active-site copper reduction promotes substrate binding of fungal lytic polysaccharide monooxygenase and reduces stability J Biol Chem 293, 1676–1687

71. Sørensen, H. V., Montserrat-Canals, M., Prévost, S., Vaaje-Kolstad, G., Bjerregaard-Andersen, K., Lund, R., et al. (2023) Tangled up in fibers: How a lytic polysaccharide monooxygenase binds its chitin substrate bioRxiv 10.1101/2023.09.21.558757

72. Suzuki, K., Suzuki, M., Taiyoji, M., Nikaidou, N., and Watanabe, T. (1998) Chitin binding protein (CBP21) in the culture supernatant of Serratia marcescens 2170 Biosci Biotechnol Biochem 62, 128–135

