## Supplementary information for "Functional characterization of multi-domain LPMOs from marine *Vibrio* species reveals modulation of enzyme activity by domain-domain interactions"

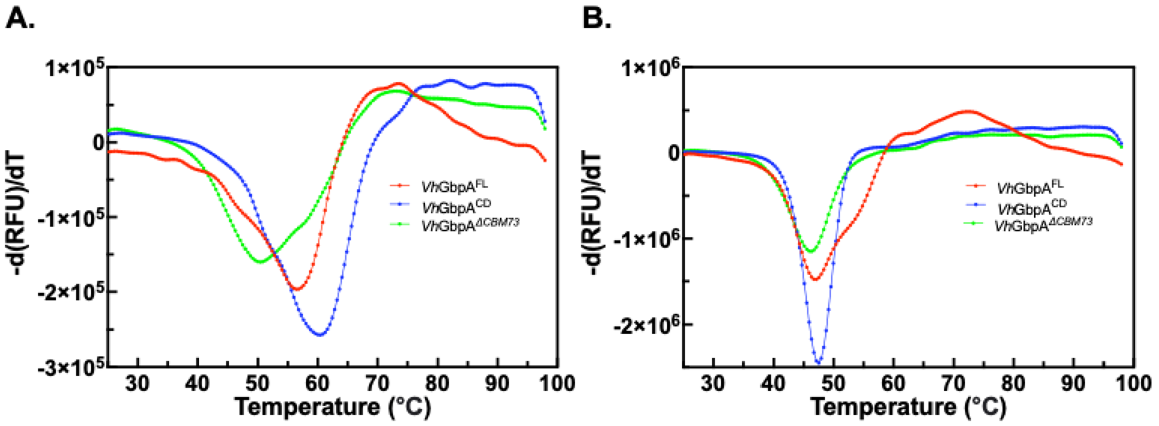


| | Apparent $T_m$ (°C ) | |
|---|---|---|
| | Without 5 mM EDTA | With 5 mM EDTA |
| *Vh*GbpA$^{FL}$ | 56.4 ± 0.0 | 47.1 ± 0.1 |
| *Vh*GbpA$^{CD}$ | 60.4 ± 0.1 | 47.5 ± 0.1 |
| *Vh*LGbpA$^{\Delta CBM73}$ | 50.4 ± 0.0 | 46.3 ± 0.0 |

**Figure S1. Thermal unfolding of *Vh*GbpA variants.** The reaction mixtures contained 5 µM enzyme and SYPRO Orange dye, incubated either in the absence (A) or presence (B) of 5 mM EDTA in 20 mM Tris-HCl buffer (pH 7.5). The temperature was gradually increased from 25°C to 99°C over 75 minutes. Each experiment was carried out in triplicate ($n = 3$) to ensure reproducibility. The table below the graphs present the apparent melting temperatures ($T_m$) derived from the unfolding curves.

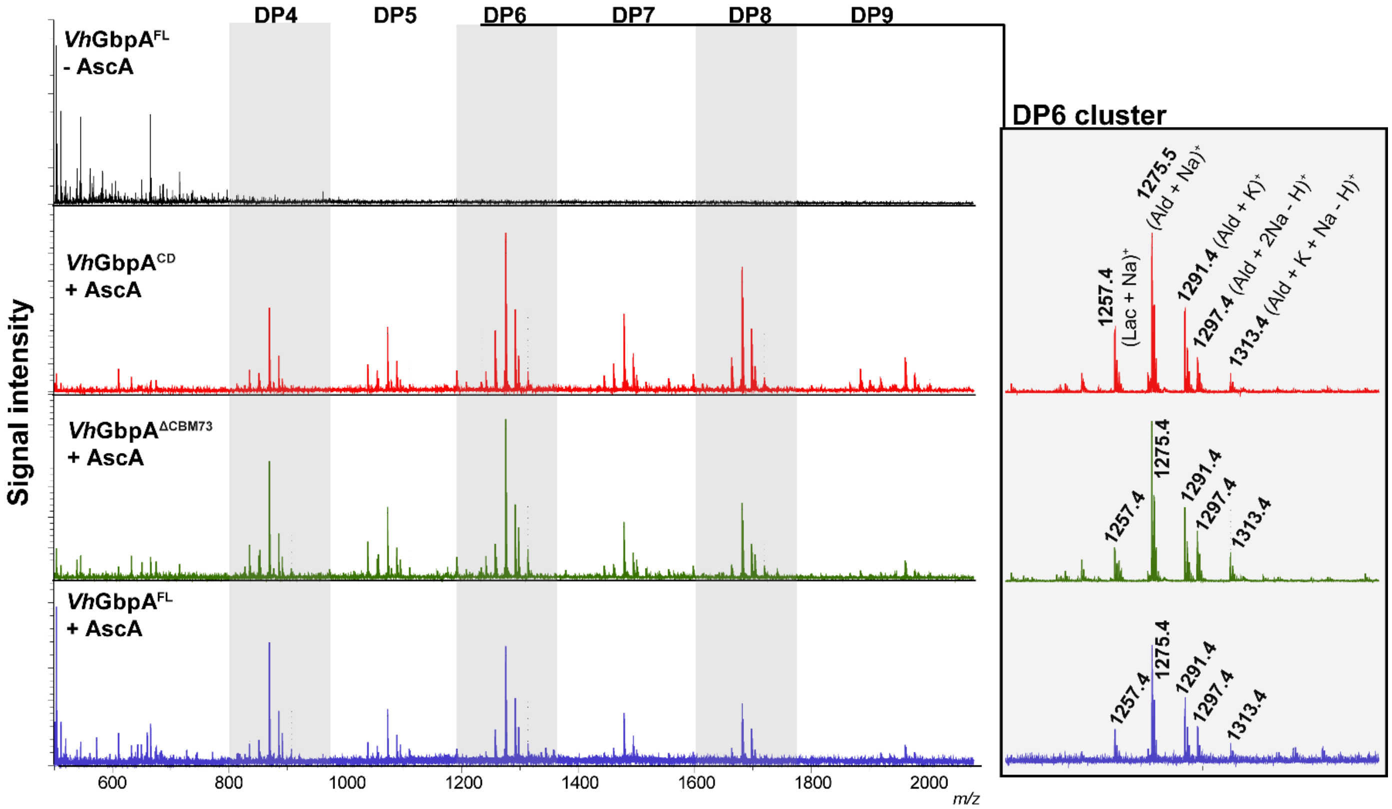


**Figure S2. Oxidized products generated by *Vh*GbpA variants reacting with chitin.** Reactions were performed 0.5µM LPMO and 10 g.L$^{-1}$ β-chitin in 20 mM Tris–HCl buffer (pH 7.5), fueled by 1 mM AscA at 30°C with shaking at 1000 rpm for 24 hours. The reactions were stopped by filtration using a MultiScreen™ 96-well filter plate operated with a Millipore vacuum manifold. Product mixtures were subsequently analyzed using MALDI-TOF MS. The left panel shows raw spectra of the reaction mixtures (blue, *Vh*GbpA$^{FL}$; red, *Vh*GbpA$^{CD}$; green, *Vh*GbpA$^{\Delta CBM73}$), whereas the right panel provides a zoomed-in view of the signals corresponding to hexameric products, with annotation of multiple signals. A control reaction with *Vh*GbpA$^{FL}$ but without AscA was also included, represented by the top black spectrum in the left panel.

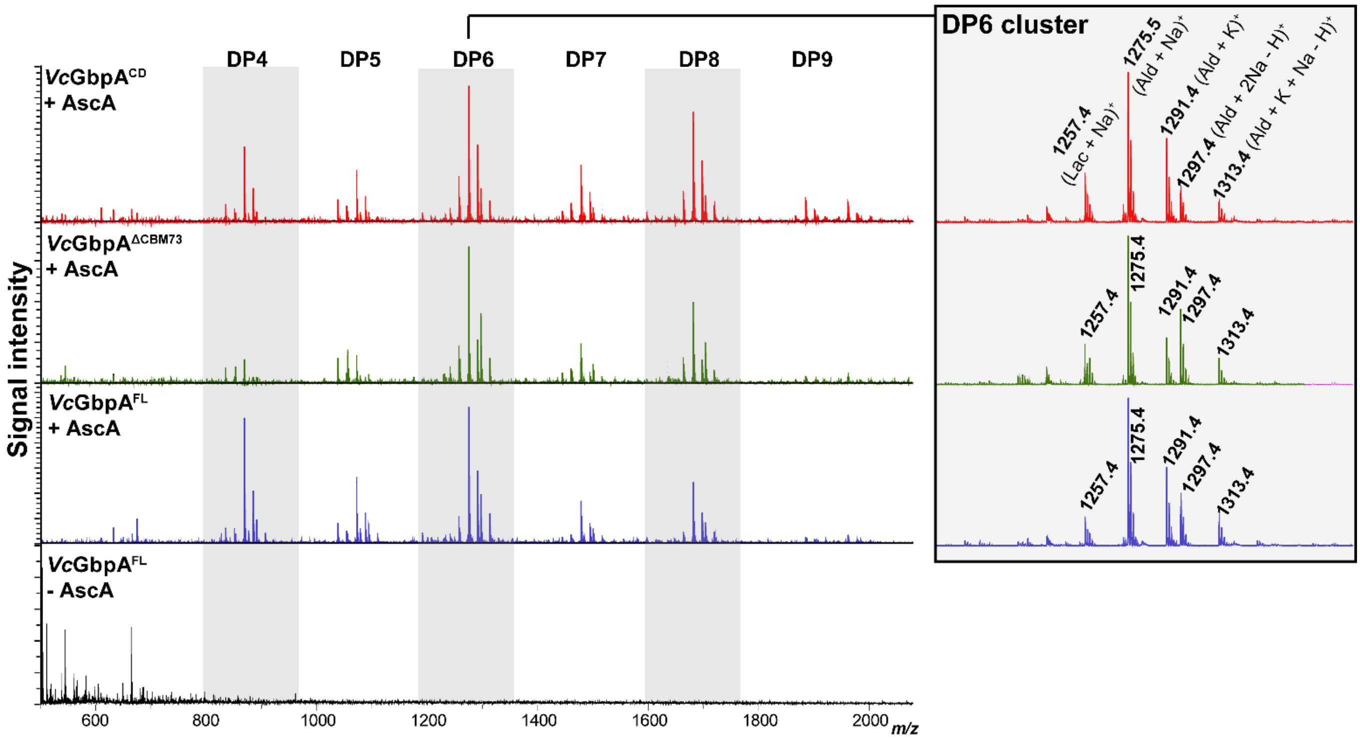


**Figure S3. Oxidized products generated by *Vc*GbpA variants reacting with chitin.** Reactions were performed 0.5µM LPMO and 10 g.L$^{-1}$ β-chitin in 20 mM Tris–HCl buffer (pH 7.5), fueled by 1 mM AscA at 30°C with shaking at 1 000 rpm for 24 hours. The reactions were stopped by filtration using a MultiScreen™ 96-well filter plate operated with a Millipore vacuum manifold. Product mixtures were subsequently analyzed using MALDI-TOF MS. The left panel shows raw spectra of the reaction mixtures(blue, *Vc*GbpA$^{FL}$; red, *Vc*GbpA$^{CD}$; green, *Vc*GbpA$^{\Delta CBM73}$), whereas the right panel provides a zoomed-in view of the signals corresponding to hexameric products, with annotation of multiple signals. A control reaction with *Vc*GbpA$^{FL}$ but without AscA was also included, represented by the bottom black spectrum in the left panel.

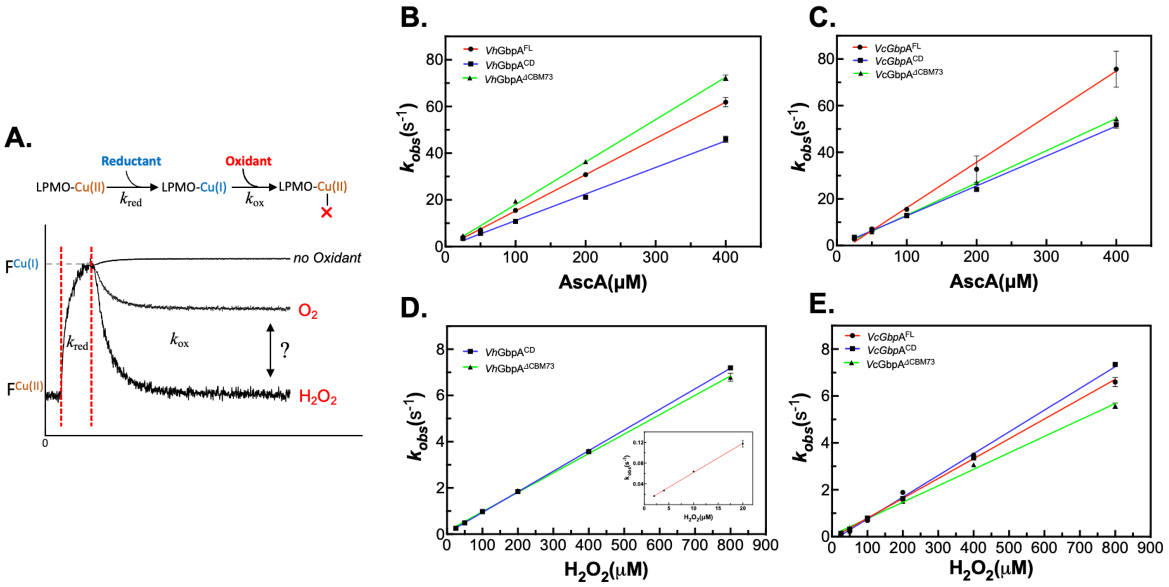


**Figure S4. Reduction and reoxidation rates. A**. Illustration of LPMO reduction and reoxidation reactions and expected related changes in intrinsic fluorescence. Upon reduction, the intrinsic fluorescence (F) of the LPMO increases, while it decreases upon subsequent reoxidation by $O_2$ (slower) or $H_2O_2$ (faster; see main text). **B-E.** Pseudo-first-order rate constants, $k_{obs}$, obtained in 20 mM Tris-HCl, pH 7.5, at 25°C, are plotted as a function of the concentration of AscA (reduction; panels **B, C**) or $H_2O_2$ (reoxidation; panels **D, E**). For *Vh*GbpA$^{FL}$, reoxidation could not be determined using stopped-flow fluorometry and was therefore measured by fluorescence spectroscopy instead (inset in panel **D**). Lower (2-20 μM) $H_2O_2$ concentrations were used because the manual setup does not allow immediate measurement after mixing, unlike the rapid mixing capability of stopped-flow, which is essential when using high $H_2O_2$ concentrations due to the fast reoxidation reaction. Each experiment was performed in triplicates and the error bars show ± S.D. (n = 3). The second order rate-constants derived from these plots are presented in **Table 1** in the main text.

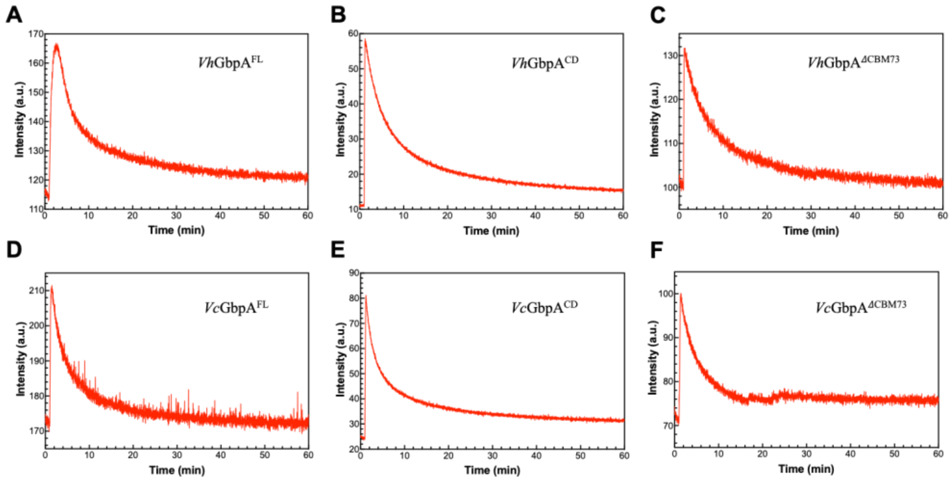


| | $k_{O2}$ (min$^{-1}$) |
|---|---|
| *Vh*GbpA$^{FL}$ | 0.049 |
| *Vc*GbpA$^{FL}$ | 0.066 |
| *Vh*GbpA$^{CD}$ | 0.055 |
| *Vc*GbpA$^{CD}$ | 0.069 |
| *Vh*GbpA$^{\Delta CBM73}$ | 0.058 |
| *Vc*GbpA$^{\Delta CBM73}$ | 0.052 |

**Figure S5. Reoxidation by $O_2$.** Reaction mixtures contained 2 µM enzyme in 20 mM Tris-HCl, pH 7.5, and were kept at 25 °C. Reduction was achieved by addition of 2 µM L-cysteine after approximately 1 minute and is reflected in an increase in the fluorescent intensity ($\lambda_{Ex/Em}$ = 280/342 nm). The subsequent decay curves reflect reoxidation by ambient molecular oxygen (approximately 250 mM). Reoxidation rates were derived by fitting the curves to $y = a + b_1 {*} e^{-k_{H2O2} {*} t} + b_2 {*} e^{-k_{O2} {*} t}$ and the resulting values are reported in the Table to the right of the Figure.

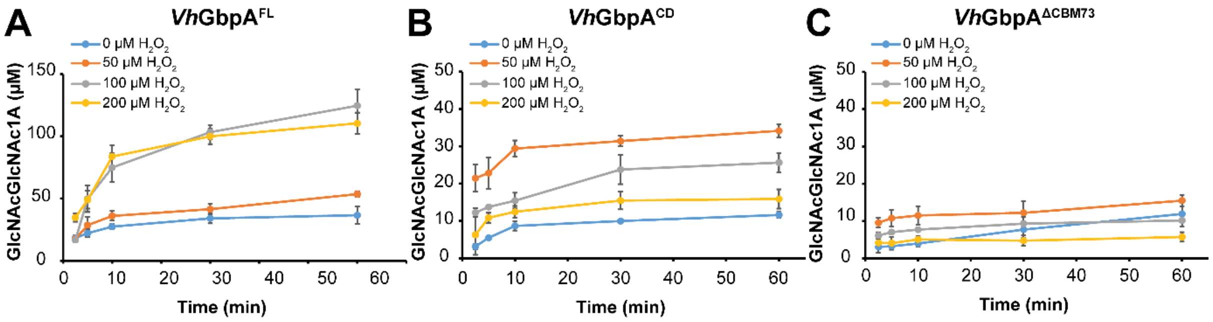


**Figure S6. $H_2O_2$-driven degradation of chitin by *Vc*GbpA variants.** The graphs show time courses for the formation of soluble oxidized products in reactions containing 0.5 μM *Vc*GbpA$^{FL}$ (**A**), *Vc*GbpA$^{CD}$ (**B**) or *Vc*GbpA$^{\Delta CBM73}$(**C**), 10 g.L$^{-1}$ β-chitin in 20 mM Tris-HCl, pH 7.5. After pre-incubating these mixtures at 30°C with shaking at 1,000 rpm for 30 mins, the reactions were initiated by sequentially adding $H_2O_2$ (to the indicated final concentrations) and, lastly, 0.1 mM AscA to start the reaction. Soluble oxidized products were quantified as described in the legend of **Fig. 3** (only chitobionic acid is shown). All reactions were performed in triplicates and the error bars show ± S.D. (n = 3). Note that product formation in the reaction with 0 mM $H_2O_2$ reflects reductant-driven enzyme activity; in these reactions $H_2O_2$ is generated slowly, *in situ*.

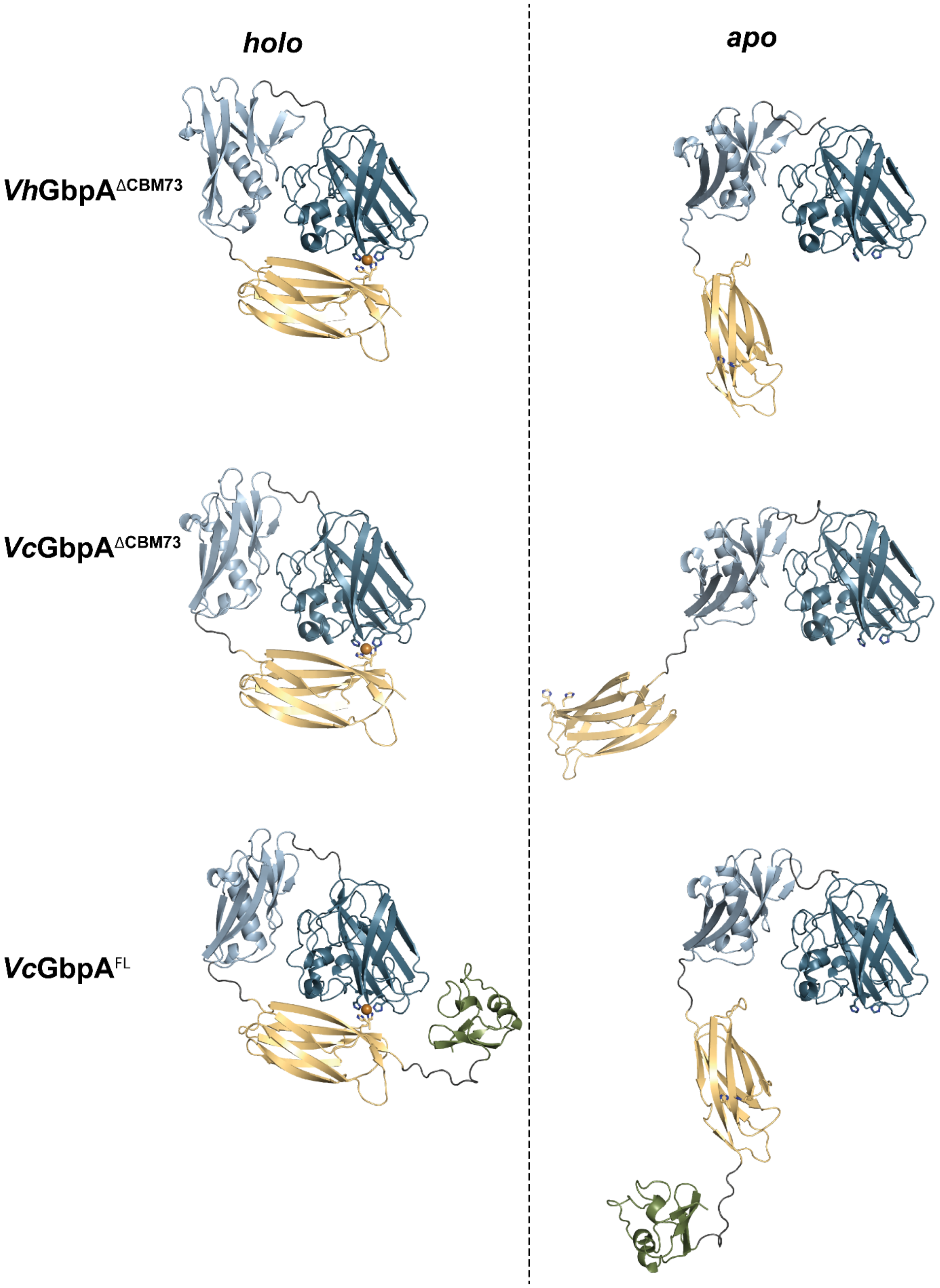


**Figure S7. Cartoon representations of *apo*- and *holo*-GbpA structures predicted using AlphaFold3.** The predicted copper-loaded structures of *Vh*GbpA$^{\Delta CBM73}$, *Vc*GbpA$^{\Delta CBM73}$, and *Vc*GbpA$^{FL}$ are shown on the left, all showing an interaction between the GbpA3 domain and the catalytic site of the AA10 domain. This interaction occurs independently of the presence of the CBM73 domain. In contrast, the predicted *apo* structures (lacking the copper cofactor) adopt more elongated conformations, and no interaction between the GbpA3 and AA10 domains is predicted.

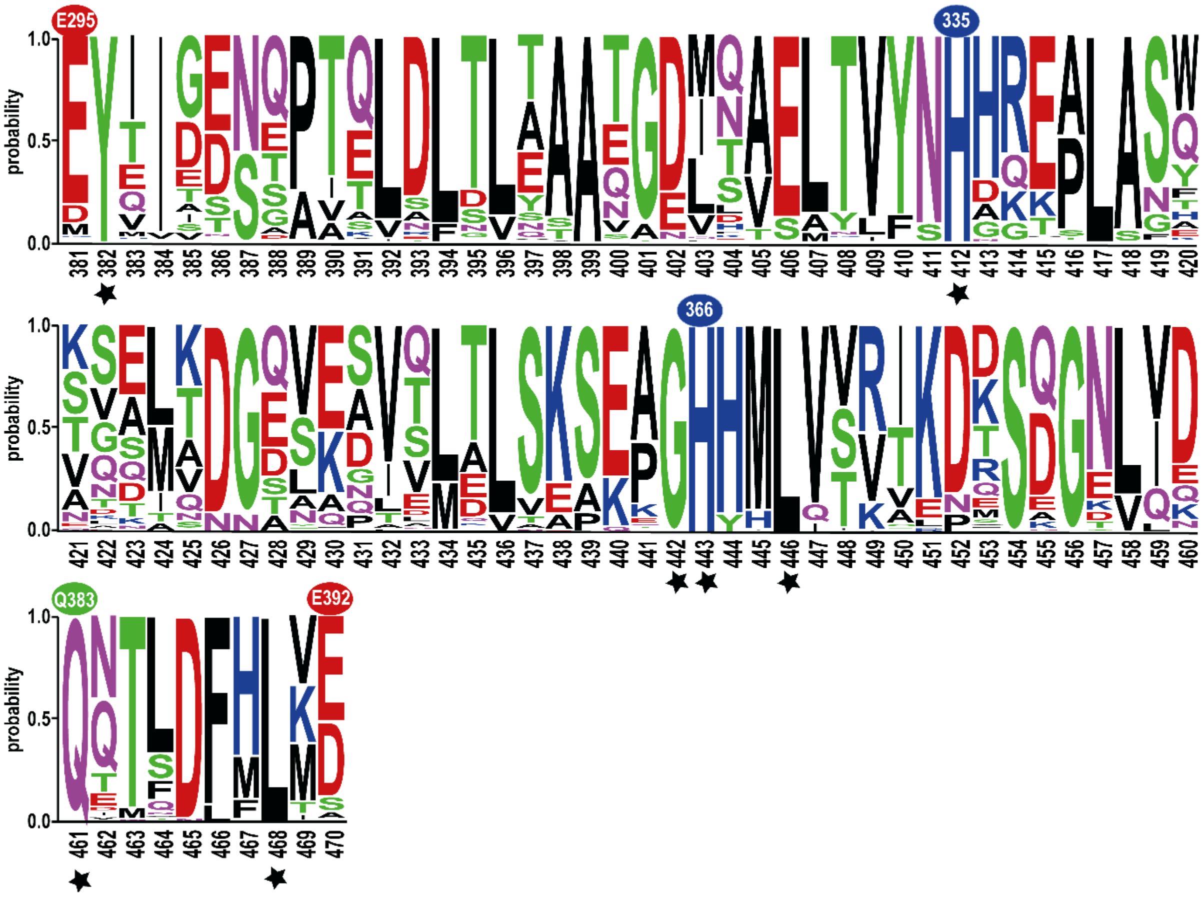


**Figure S8**. **Amino acid frequency per position based on a multiple sequence alignment of 165 sequences encoding GbpA3 domains associated with LPMOs.** The x-axis shows the MSA position numbering in black. White numbers on red or blue backgrounds highlight specific residues in GbpA3 (numbered according to *Vh*GbpA3): Glu295 and Glu392 mark the N- and C-terminal boundaries of the domain, while His335 and His366 are conserved histidines predicted to coordinate the active site copper in the AlphaFold3 model. A green background marks Gln383, which is discussed in the main text as a potential interaction point with Arg20 and Glu39 in the catalytic domain. Note that residue numbering in *Vh*GbpA excludes the signal peptide; thus, the first histidine in the mature protein is referred to as His1. Black stars beneath the x-axis indicate positions that are 100% conserved across all 165 sequences. The graph was generated using WebLogo (1).

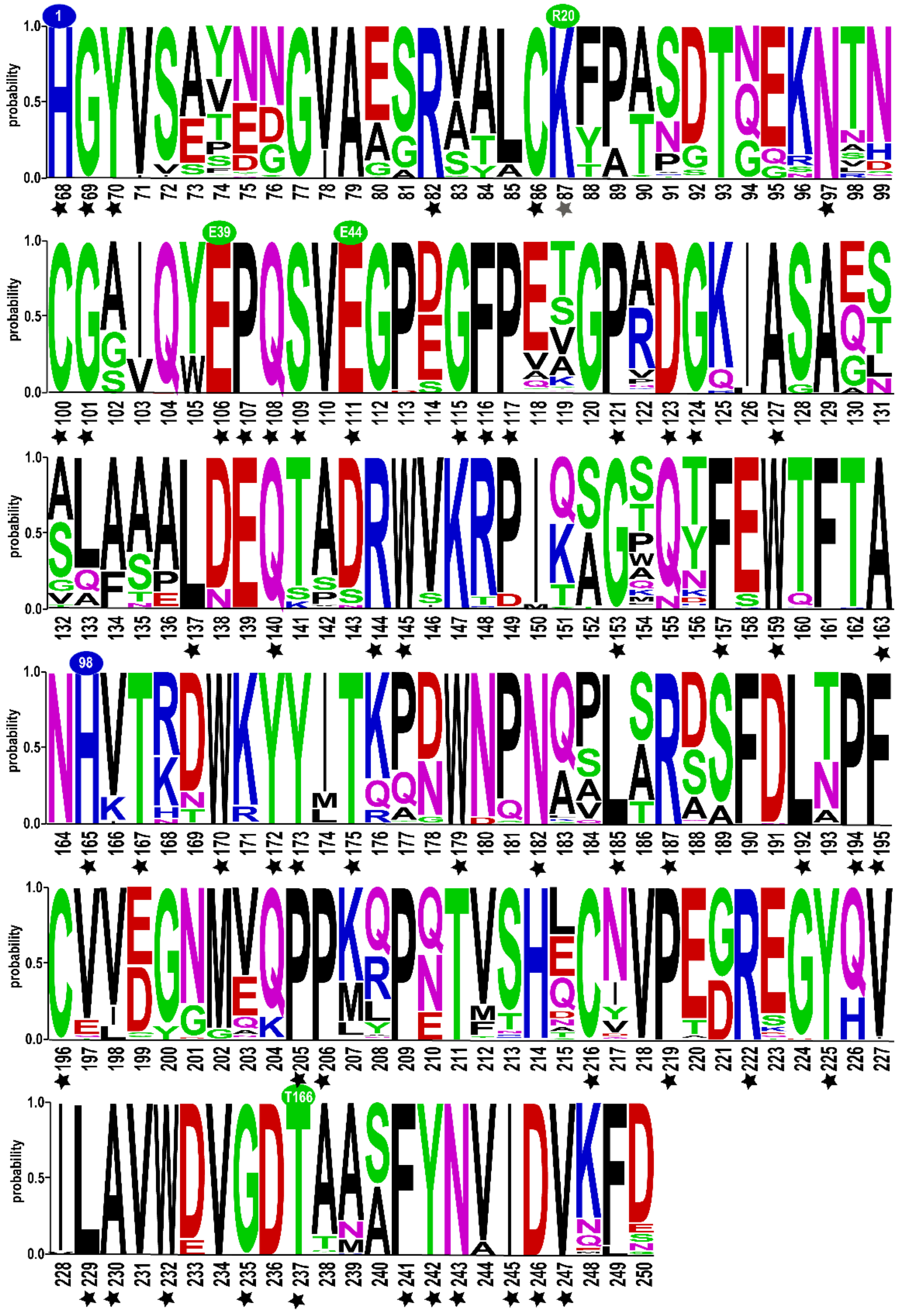


**Figure S9. Amino acid frequency per position in the AA10 domains of 165 GbpA-like sequences.** The x-axis shows the MSA position numbering in black. White numbers on blue backgrounds highlight the copper-binding histidines, His1 and His98, while white numbers on a green background indicate residues potentially involved in interactions between the AA10 and GbpA3 domains, as discussed in the main text and illustrated in **Fig. S10**. Black stars beneath the x-axis indicate positions that are fully conserved (100%) across all 165 sequences. The grey star below R20 highlights an exception found in *Vh*GbpA, where this position is occupied by an arginine instead of the lysine present in the remaining 164 sequences. Despite this substitution, the side-chain properties and positive charge are retained. The overall pairwise sequence identity across the full-length proteins (comprising all four domains) ranges from 45.4% to 99.2%, with an average pairwise identity of 51.0 ± 11.7%. The graph was generated using WebLogo (1).

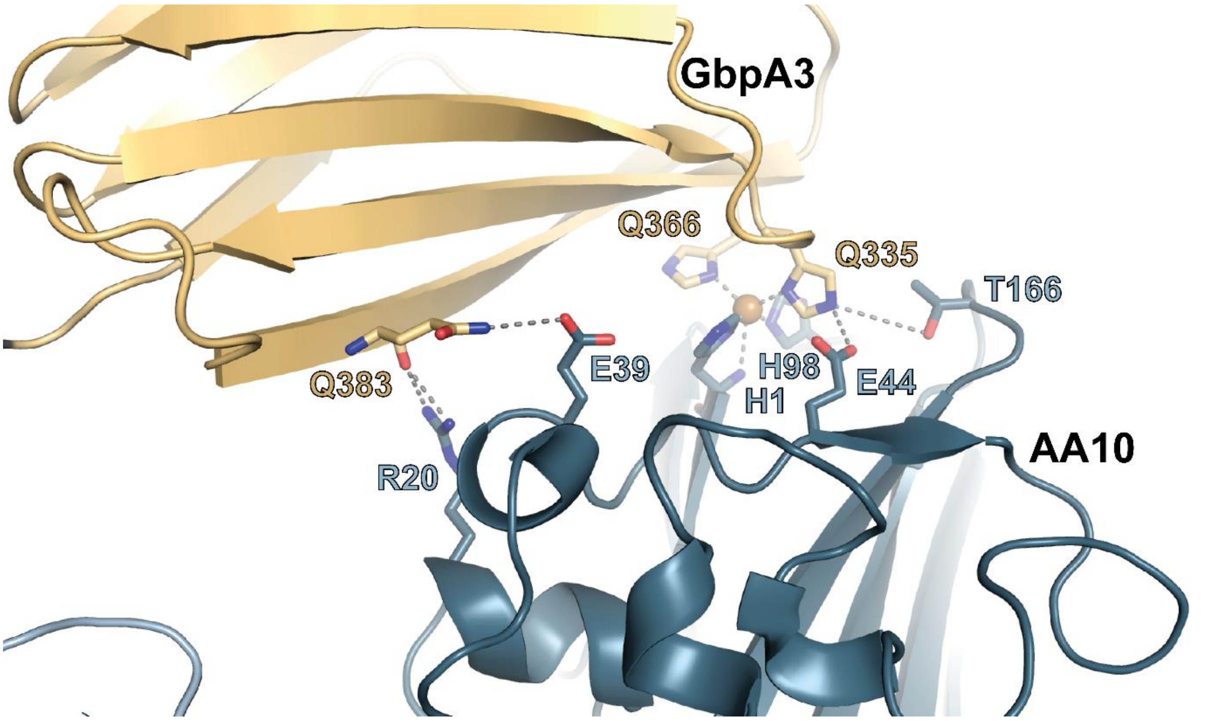


**Figure S10. Possible interactions between the GbpA3 domain and the AA10 domain in *holo*-*Vh*GbpA.** The figure shows the AlphaFold-predicted structure of *Vh*GbpA$^{FL}$, with the AA10 domain colored blue and the GbpA3 domain colored in light sand. Highlighted residues represent potential additional contact points beyond the two copper-coordinating histidines (His335 and His366). These residues are conserved among the 165 GbpA-like sequences used to generate the WebLogos shown in **Figs. S8** and **S9**.

**Table S1. The oxidase rate of LPMO variants.** Reaction mixtures containing 0.1mM Amplex Red, 5 U.mL$^{-1}$ HRP, and 1 μM enzyme in 20 mM Tris-HCl buffer (pH 7.5) were pre-incubated at 30°C for 5 mins. The reaction was initiated by adding 1mM AscA (final concentration). The solutions were mixed by shaking the plate at 600 rpm for 30 s, and absorption at 563 nm was measured every 30 s for 60 mins. A control reaction lacking the enzyme was included. Oxidase rates were derived from linear progress curves and were corrected for the signal obtained in the control reaction lacking the LPMO. The values represent the mean ± standard deviation for three independent replicates.

| | **$H_2O_2$ production rates (nM.s$^{-1}$)** | | |
|---|---|---|---|
| | Full-length | AA10 CD | ΔCBM73 |
| *Vh*GbpA | 3.4 ± 0.1 | 11.0 ± 0.2 | 5.9 ± 0.1 |
| *Vc*GbpA | 3.0 ± 0.1 | 8.8 ± 0.2 | 2.5 ± 0.1 |

### References

1. Crooks, G. E., Hon, G., Chandonia, J. M., and Brenner, S. E. (2004) WebLogo: a sequence logo generator *Genome Res* **14**, 1188-1190
